# Knowledge-informed multimodal cfDNA analysis improves sensitivity and generalization in cancer detection

**DOI:** 10.1101/2025.10.20.683167

**Authors:** Antonio De Falco, Piera Grisolia, Raffaella Giuffrida, Clara Iannarone, Cinzia Graziano, Marianna Scrima, Alessia Maria Cossu, Rossella Tufano, Maria Vanessa Yow, Chloe Marissa Brown, Palak Bajaj, Marco Bocchetti, Pier Vitale Nuzzo, Floriana Morgillo, Francesco Caraglia, Maria Carminia Delle Corte, Gaetano Di Guida, Teresa Troiani, Fortunato Ciardiello, Maria Rosaria Rizzo, Alfonso Fiorelli, Noemi Maria Giorgiano, Davide Arcaniolo, Giampiero Della Rosa, Marco De Sio, Alessia Covre, Anna Maria Di Giacomo, Luana Calabrò, Paolo Fontana, Marzia Mare, Ola Landgren, Damian J. Green, Alex Lesokhin, David G. Coffey, Nipun Merchant, Jashodeep Datta, Stefano Forte, Michele Maio, Michele Caraglia, Michele Ceccarelli

## Abstract

Liquid biopsy offers a minimally invasive opportunity to detect and monitor cancers through analysis of cell-free DNA (cfDNA). However, current approaches face challenges of limited sensitivity at low tumor fractions, technical variability, and poor generalization across cohorts. Tumor-informed targeted methods offer high specificity but suffer from low sensitivity due to random sampling, tumor evolution and adaptation (including resistance mechanisms), and other sources of heterogeneity. Conversely, tumor-naive genome-wide methods can increase sensitivity but often sacrifice specificity, particularly at low tumor fractions. We developed Fragmentomics Analysis for Tumor Evaluation with AI (Fate-AI), a multimodal framework that integrates fragmentomic and methylation-derived features from low-pass whole-genome sequencing (LPWGS) and cell-free methylated DNA immunoprecipitation and high-throughput sequencing (cfMeDIP-seq). It employs a knowledge-informed strategy to select recurrently altered genomic regions and tissue-specific methylation loci to combine the advantages of tumor-naive approaches with the specificity of tumor-informed approaches. This approach derives robust per-sample normalized features that mitigate batch effects and enhance cross-cohort reproducibility. We evaluated Fate-AI on a total of 1,219 plasma samples spanning ten cancer types and healthy controls from multiple laboratories and sequencing centers, including 432 newly profiled cases (280 with both cfMeDIP-seq and LPWGS) together with 787 samples from four independent public datasets. Fate-AI achieved superior sensitivity and specificity compared to state-of-the-art methods, detecting tumor-derived signals at fractions as low as 10^−5^ in experimental dilutions. Fate-AI scores correlated with disease stage and tracked longitudinal progression, anticipating relapse months before clinical progression. Furthermore, Fate-AI enabled tissue-of-origin classification, with AUCs ranging from 0.84 to 0.97 across six cancer types. Collectively, our results demonstrate that Fate-AI provides a sensitive, generalizable, and clinically actionable platform for early detection, minimal residual disease monitoring, and tissue-of-origin classification, supporting its potential as a liquid biopsy framework in precision oncology.

## Introduction

Liquid biopsy, through the non-invasive analysis of circulating tumor DNA (ctDNA), the fraction of cell-free DNA (cfDNA) derived from tumor cells, offers transformative opportunities for cancer management. Applications span early detection, tissue-of-origin classification, genotyping, and minimal residual disease (MRD) monitoring[1, 2]. The targeted ultra-deep sequencing of specific genomic regions defined by tumor-informed bespoke panels or recurrent driven events has been used to overcome the sparse nature and the low fraction of ctDNA [3, 4, 5, 6, 7, 8]. However, the sensitivity of such assays remains limited because, despite the use of ultra-deep sequencing, somatic mutations can be undetectable if they are absent from the finite plasma volume obtained during standard sample collection, creating an intrinsic sensitivity limitation [9, 10, 11, 12]. Current approaches are effective in patients with advanced malignancies, but have limited efficacy at early stages. Moreover, tumor-informed panels also have the intrinsic limitation of not accounting for tumor evolution and acquired resistance mechanisms. Recently, genome-wide tumor-informed approaches have emerged to overcome these limitations by integrating thousands of single-nucleotide variants and copy number alterations across the entire genome [13, 14, 15]. Other tumor-naive approaches exploit the fact that cell-free DNA coverage across the genome is uneven, and nucleosome spacing highly correlates with epigenetic footprints of the tissues from which the cfDNA is released [16]. Since epigenetic states are highly tissue-specific, such footprints can be used to deconvolve the cfDNA and identify the organs or tissues releasing the DNA fragments observed in circulation. This concept has been used to develop innovative liquid biopsy approaches in prenatal testing, oncology, and monitoring of organ transplant recipients [17]. The distribution of fragmentomics features such as the ratio between long and short fragments [18], end-motif [19], read-depth of transcription factor binding sites [20], and other multi-modal approaches [21] have been used for tumor-naive cancer detection. Other epigenetic footprints characterizing ctDNA are associated with methylation patterns that can be detected by immunoprecipitation of methylated DNA [9] and can accurately detect renal cell carcinoma [22], discriminate between patients with localized and metastatic prostate cancers [23], classify intracranial tumors [24], and even contribute to the identification of cancer subtypes [25].

The size distribution of DNA fragments from ctDNA often differs from that in normal cells ([26, 19, 27]). This variance might arise from specific cell death mechanisms, unusual nuclease activity, particularly involving the endonuclease enzyme DNASE1L3 [28], and the genomic instabilities associated with cancer that can lead to the production of more fragmented cfDNA. It has been definitely shown that the fragmentation pattern reflects the organization of nucleosomes [16, 29, 30]. However, fragment distribution in sequencing data can be influenced by technical variations, often referred to as batch effects or sequencing effects. These effects arise due to differences in sample preparation, sequencing platforms, reagent batches, or processing pipelines, potentially leading to biases in fragment size distribution across different cohorts [31]. To overcome these challenges, we reasoned that a potential approach to reduce the variability could be a per-sample normalization that differentiates the same features in multiple genomic regions of the same sample. This could facilitate comparisons among different samples and batches. We, therefore, implemented Fragmentomics Analysis for Tumor Evaluation (Fate-AI). Our method adopts a knowledge-informed approach, where regions known to be enriched for genomics and epigenomic alterations are considered to extract fragmentation-based features. We use regions of frequently amplified and deleted regions and regions frequently methylated in the specific tumor. For this purpose, we integrate the features extracted for LPWGS and cfMeDIP-seq [9]. We demonstrate that Fate-AI achieves higher sensitivity and specificity than state-of-the-art approaches, particularly when different datasets are used for external cross-validation.

## Results

### Fragmentation patterns differ across studies and conditions

We evaluated the distribution of circulating DNA fragment lengths in whole-genome sequencing data from pre-treatment cancer patients and healthy controls, using both our cohort (n=315) and published datasets (n=419). As in many previously reported studies [17, 18, 32, 30], a distinct shift in fragment size distributions was observed between the cancer and control. In cancer patients, fragments were enriched around 145 base pairs (bp), corresponding to DNA wrapped around the nucleosome core particle (without linker DNA). In contrast, healthy individuals showed a higher proportion of fragments extending to 197 bp, which reflects DNA associated with the chromatosome, consisting of the nucleosome plus the linker histone H1 and its 50 bp of protected DNA (**Figure 1A**). This difference confirms that chromatin organization and nuclease cleavage patterns in cfDNA differ between tumor-derived and normal sources. Interestingly, the extent of the shift toward shorter fragments in cancer patients correlated with tumor content, with cohorts characterized by a higher tumor fraction in cfDNA showing a more pronounced effect. This is particularly evident for the samples in the samples from the cohorts reported in [25] and [21] having a greater tumor content and a consequent greater shift between the fragment length distribution between cancer and healthy cases (**Figure 1A** and **Supplementary Figure S1A**). To mitigate potential bias, we selected samples spanning multiple cancer types but characterized by relatively lower tumor fractions, representing particularly challenging cases (**Supplementary Figure S1A**). Unfortunately, the phenotypic differences between pathological cases and healthy individuals are not the only source of difference in fragment length distribution. When comparing healthy samples across multiple studies, a clear batch effect emerged, with differences in fragment length distributions reflecting the influence of study design and experimental protocols (**Figure 1B**). A similar effect was evident across different cohorts of the same cancer type, with each cohort showing distinct distributional patterns (**Supplementary Figure S1B**). This shift must be considered when training and evaluating the generalization performance of tumor-naive cfDNA analysis methods. This discrepancy significantly challenges the distinction between the biological effects and technical variations. This issue is one of the main obstacles to the generalization of several predictive models to multiple cohorts [31]. Therefore, a critical need in the field of liquid biopsy is to characterize the cfDNA epigenetic profiles with features that remain consistent between cohorts while minimizing the impact of batch effects and technical variability. This could improve the reproducibility of studies and the generalization of predictive models.

**Figure 1.**
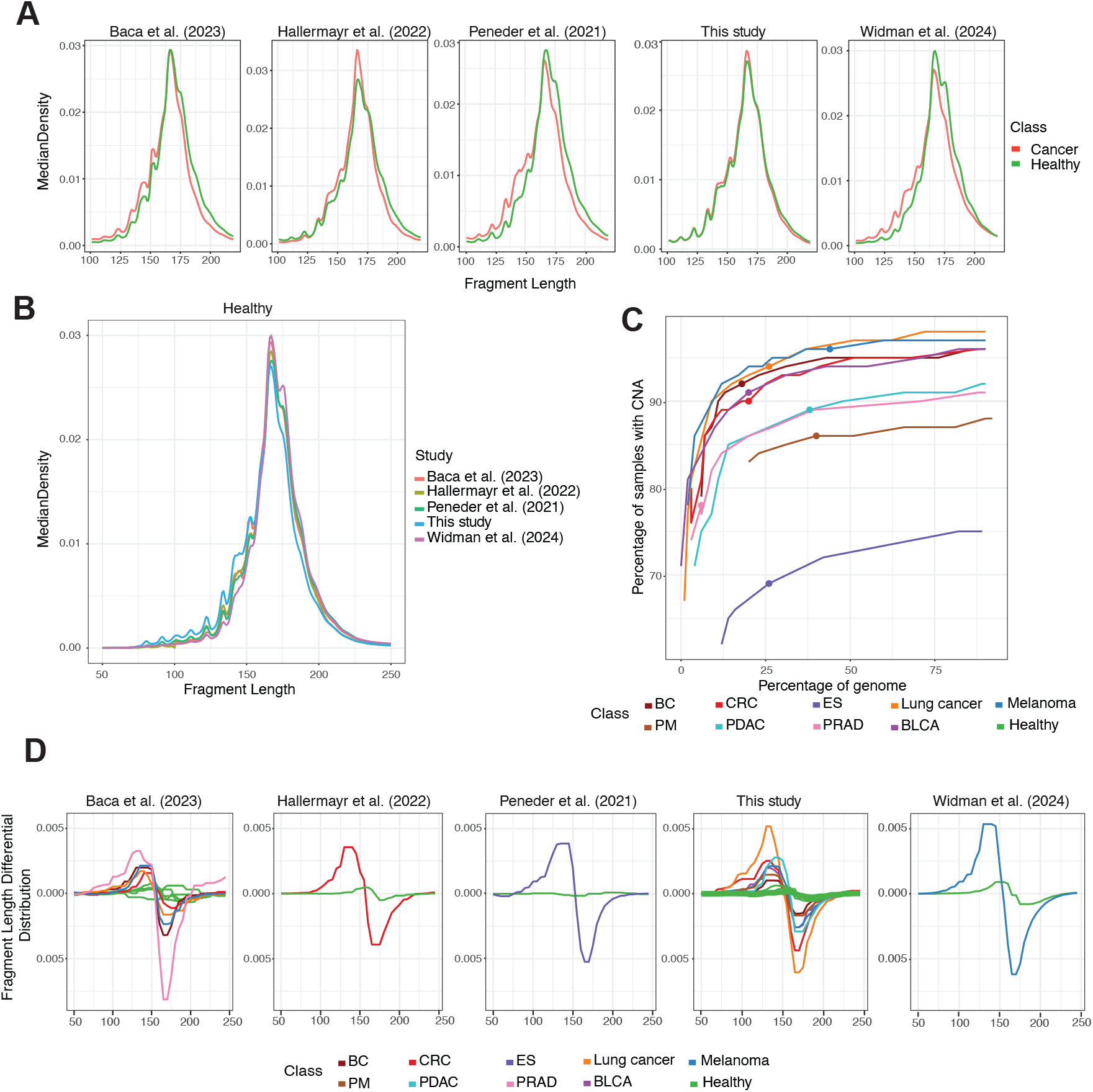
Fragment length and copy number alteration (CNA) frequencies across multiple datasets. **A**, Density of cfDNA fragment lengths for cancer samples and healthy controls. Data from Baca et al. [25] (healthy n=33, cancer n=122), Hallermayr et al. [35], (healthy n=61, cancer n=43), Peneder et al. [21] (healthy n=21, cancer n=66), this study (healthy n=69, cancer n=246), Widman et al. [15] (healthy n=35, cancer n=38). **B**, Density of cfDNA fragment lengths from 219 healthy controls across different datasets [25, 35, 21, 15]. **C**, The percentage of samples harboring at least one CNA as a function of the percentage of the genome affected, varying the recurrency thresholds from 2.5% to 50% in steps of 2.5, stratified by cancer type. Points represent the selected thresholds for all the reported experiments. Data from the Progenetix database [34]. **D**, Fragment Length Differential Distribution for each cancer type across datasets [25, 35, 21, 15]. Abbreviations: breast cancer (BC), colorectal carcinoma (CRC), Ewing sarcoma (ES), lung cancer, pleural mesothelioma (PM), pancreatic ductal adenocarcinoma (PDAC), urothelial bladder carcinoma (BLCA).

Large copy number alterations (CNAs), gains and losses of chromosomal regions, are one of the main hallmarks of cancer [33]. We analyzed CNAs across multiple cancer types by considering regions of recurrent CNA events and quantifying the proportion of samples harboring at least one CNA within these regions. To this end, we partitioned the genome into bins of 3, 000, 000 bp and estimated the fraction of the genome affected by CNAs at varying recurrence thresholds, using a dataset of 156,871 copy number profiles spanning diverse cancer types [34]. The relationship between the percentage of the genome encompassed by recurrent CNAs and the percentage of patients carrying at least one CNA, for the nine cancer types considered in this study, is shown in **Figure 1C**. The recurrence threshold, to select the analyzed regions, was varied from 2.5% to 50%, in steps of 2.5%. Decrease the threshold for selecting recurrently alternated regions, the considered portion of the genome increases, and the proportion of samples showing CNAs also rises, eventually reaching a plateau. For several types considered in this study, with a recurrent threshold of 25% we have at least 90% of cases affected by one or more CNAs, except for cancers characterized by lower chromosomal instability, such as Ewing Sarcoma (ES), pleural mesothelioma (PM), and pancreatic ductal adenocarcinoma (PDAC). We then evaluated differences in the distribution of cfDNA fragment lengths across selected genomic regions, specifically comparing frequently amplified versus frequently deleted loci, in the data generated in this study and public datasets. We observed substantially greater variability in fragment length distributions across CNA regions in cancer patients, whereas healthy controls exhibited almost flat profiles. We leveraged these differences in fragment length distributions to define a per-sample *Fragment Differential Distribution* (FDD). In cancer samples, FDD displayed pronounced, opposite-phase modulation patterns: a sharp positive deviation peaking at approximately 140 bp and a steep negative deviation reaching a minimum near 170 bp (**Figure 1D**). FDD was utilized to derive a set of numerical features for cancer detection, monitoring of minimal residual disease, and tissue-of-origin classification (Methods). Notably, these features were able to discriminate cancer patients from healthy controls even at very low estimated tumor fractions (*<* 0.03) (**Supplementary Figure S1C**). These results indicate that the differential distribution of fragment features can enhance detection sensitivity by capturing recurrent patterns shared between cohorts and cancer type. The use of features based on within-sample contrasts may also mitigate cohort-dependent bias because it relies on relative differences between regions within the same sample.

### Multimodal analysis of circulating free DNA features for cancer detection and classification

Based on the analysis presented above, we reasoned that a potential strategy to improve the signal for cancer detection is to contrast features from genomic regions enriched in ctDNA-derived fragments or depleted of ctDNA-derived fragments. This strategy can be applied to epigenetic footprints reflecting the nucleosome organization derived from LPWGS, as well as to tissue-specific methylation patterns derived from cfMeDIP-seq. Based on these assumptions, we developed Fragmentomics analysis for tumor evaluation with AI (Fate-AI). By adopting a knowledge-informed approach, the model contrasts the features extracted from the recurrently amplified and deleted regions of the specific tumors of interest, as well as the expected differentially methylated regions in cancer with respect to plasma (Methods). The combined set of features is used to train an elastic net model to estimate the tumor content in cfDNA (**Figure 2**).

**Figure 2.**
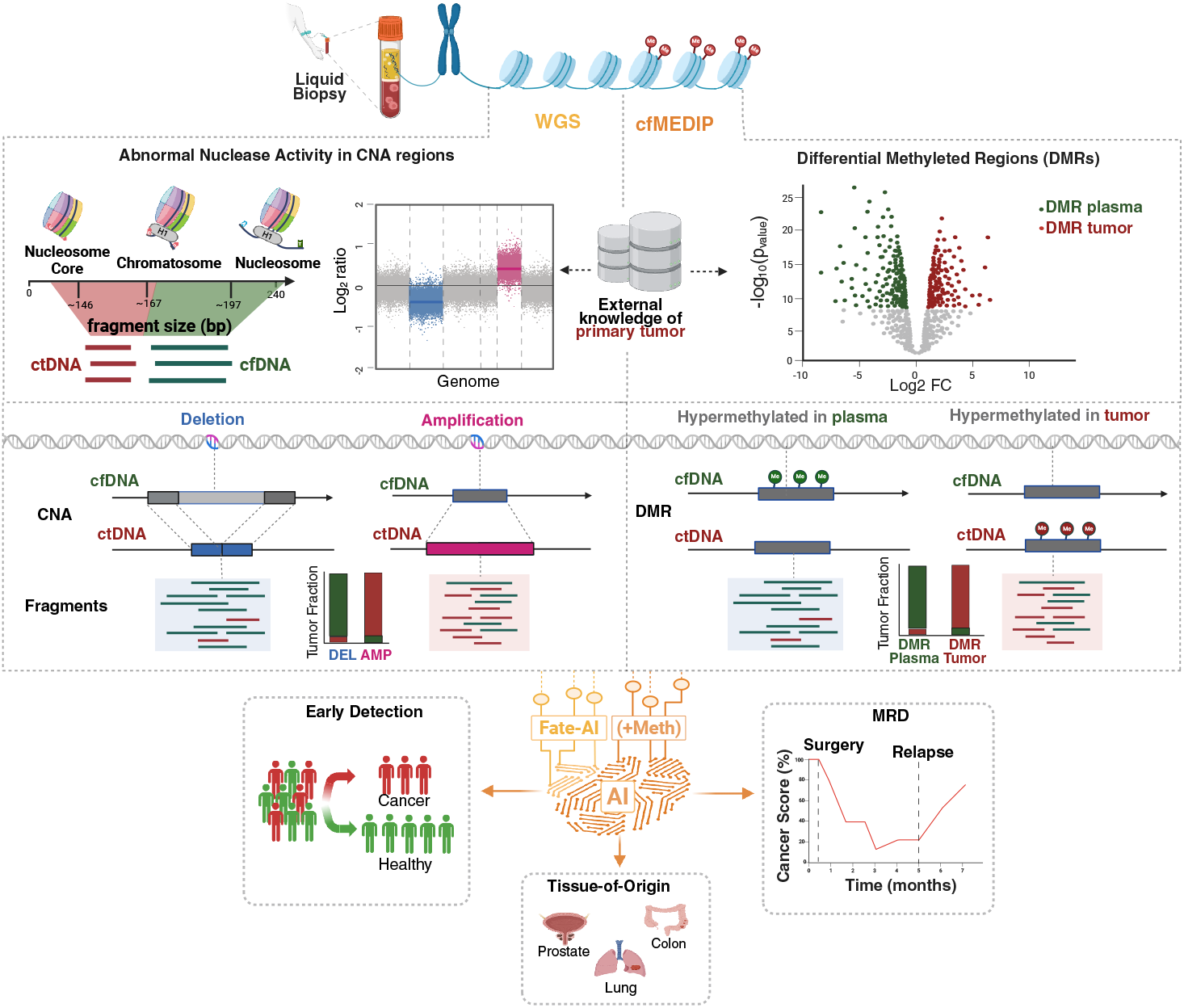
Multimodal analysis of circulating tumor DNA (ctDNA) features for cancer detection and classification. Overview of the analytical framework integrating low-pass whole-genome sequencing and cfMeDIP-seq for liquid biopsy analysis. Fate-AI calculates a per-sample Normalized Differential Distribution of multiple epigenetic footprints from LPWGS. The model assumes that, in cancer samples, amplified regions would be enriched for ctDNA-derived fragments, whereas recurrently deleted regions would predominantly contain cfDNA originating from normal cells. The differential distribution of epigenetic features has a distinct pattern between cancer and healthy samples. These features are then integrated with features from differentially methylated regions (DMRs) between plasma and tumor samples to refine tumor signal identification further. A logistic regression model for early cancer detection, minimal residual disease monitoring, and tissue-of-origin classification using artificial intelligence (AI)-driven models is then used to score the input sample.

We profiled 432 liquid biopsies, spanning eight cancer types from seven cancer centers, three laboratories, and four sequencing facilities (**Supplementary Table S1**). Our cohort included breast cancer (BC) (n=42), colorectal carcinoma (CRC) (n=50), lung cancer (n=29), melanoma (n=40), PM (n=24), urothelial bladder carcinoma (BLCA) (n=38), multiple myeloma (MM) (n=91), and PDAC (n=49), along with 69 healthy controls. For 280 cases, we generated both cfMeDIP-seq and LP-WGS profiles (**Supplementary Figure S2**). In addition, we analyzed 787 samples from four independent public datasets: CRC [25, 35] (n=228), lung cancer [25] (n=38), ES [21] (n=184), BC [25] (n=15), prostate adenocarcinoma (PRAD) [25] (n=32), melanoma [25, 15] (n=139), and healthy controls [25, 35, 15, 21] (n=150) (**Supplementary Table S2**). Among these validation cohorts, 149 samples had both cfMeDIP-seq and LPWGS available. Since we will evaluate Fate-AI on some cohorts profiled only with LPWGS, we refer to the full multimodal model as Fate-AI(+Meth) to distinguish the case where only fragmentomics features are used.

### Sensitivity of Fate-AI in Tumor-Naive Cancer Detection

We first evaluated the sensitivity of Fate-AI for tumor-naïve cancer detection across varying tumor fraction (TF) levels using *in silico* admixtures. Specifically, known proportions of tumor-derived DNA were digitally mixed with cfDNA from healthy individuals to generate synthetic datasets. Whole-genome sequencing (WGS) data from CRC, lung cancer, and melanoma samples with relatively high TF were mixed with WGS data from plasma samples from cancer-free individuals. The admixture process accounted for both the initial tumor content and the ploidy of each tumor sample to determine the dilution ratios required to achieve target TFs ranging from 10^−6^ to 10^−1^. To ensure comparability across samples, sequencing coverage was normalized to a consistent target depth. Two evaluation scenarios were considered: (i) a matched-cohort setting, models evaluated on the mixtures were trained on the same cohort from which the mixtures were generated, excluding the specific control/cancer sample pairs used to generate the mixtures, and (ii) a cross-cohort setting, in which models were trained on samples from one cohort and tested on data generated by a different study. Performance was assessed using the area under the receiver operating characteristic curve (AUC). Because AUC reflects only the relative ranking of prediction scores, we also calculated the *Delta Score*, defined as the mean difference between the average prediction scores of tumor and control samples. This metric provides an interpretable measure of class separation, facilitating the identification of diagnostic thresholds. We benchmarked Fate-AI against Delfi [18]. Since Delfi is based solely on fragment length ratios from LPWGS, we used Fate-AI without cfMeDIP-seq–derived methylation features to ensure a fair comparison. Fate-AI achieved high AUC values at TF − 10^−3^: 0.95, 0.75, and 0.82 for CRC, lung cancer, and melanoma, respectively (**Figure 3A**). Similarly, a significant Delta Score separation at TF as low as − 10^−3^, outperforming Delfi in the matched-cohort setting. In the cross-cohort setting, for CRC, we trained the model on 32 tumor and 33 normal samples from Baca et al.[25] and tested it on data from Hallermayr et al.[35]. For lung cancer and melanoma, models were trained on data from Baca et al.[25] (lung cancer n=38, healthy n=33) and Widman et al.[15] (melanoma n=38, healthy = 35), and evaluated on the same test cases of Figure 3A. Fate-AI, in this scenario, reaches consistently high accuracy in CRC with an AUC of 0.82 at TF 10^−4^, lung cancer with an of AUC 0.82 at TF 10^−3^, and melanoma with an AUC 0.9 at TF 10^−3^ (**Figure 3B**). Across all three cancer types, Fate-AI consistently demonstrated better generalization and robustness at low TFs (**Figure 3C**).

**Figure 3.**
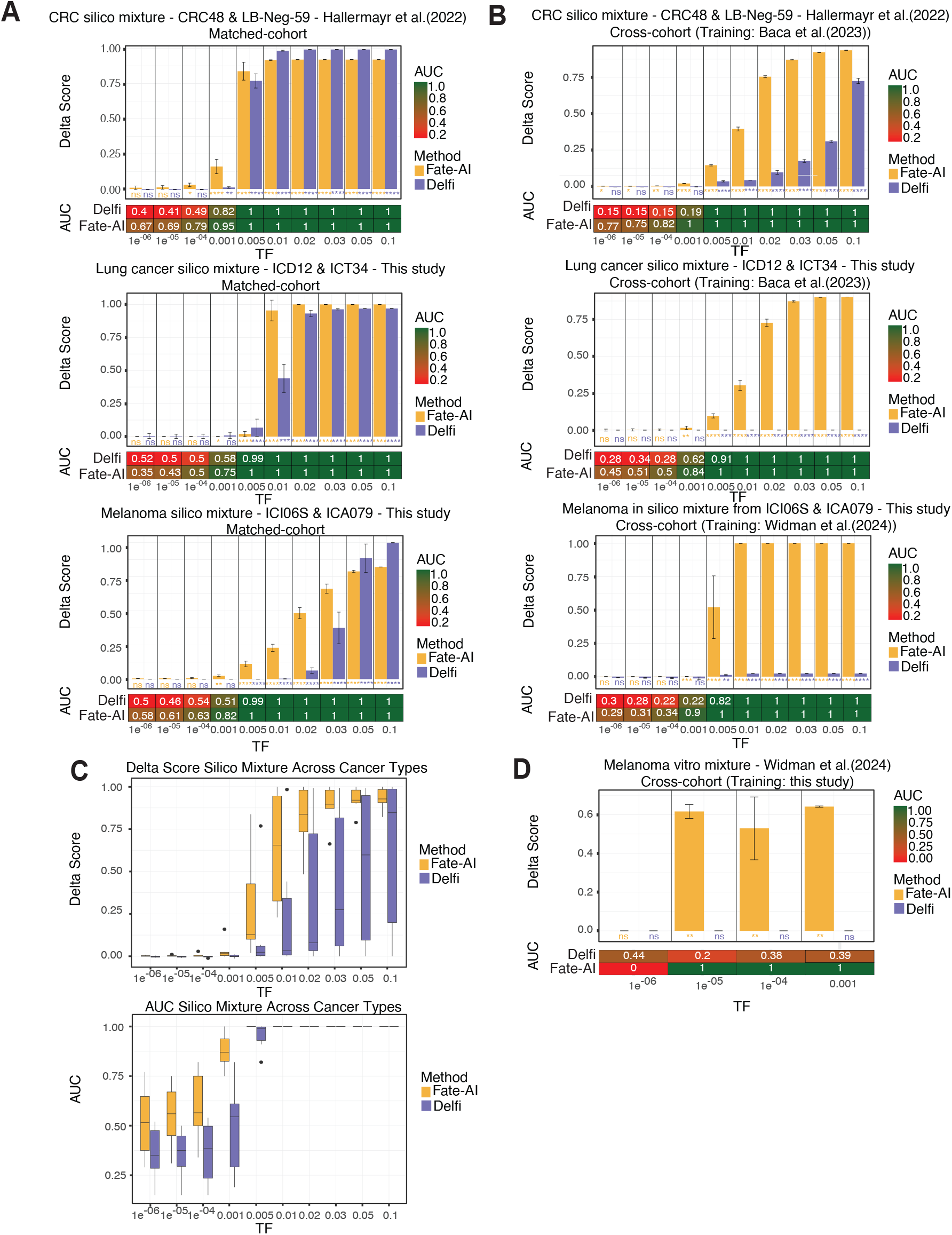
Detection sensitivity of Fate-AI on in silico and in vitro admixtures. **A**, Matched-cohort setting: models evaluated on the mixtures were trained on the same cohort from which the mixtures were generated, excluding the specific control/cancer sample pairs used to create the mixtures. For each value of synthetic TF, ten in silico mixtures of CRC, lung cancer, and melanoma were generated. For CRC, we used the sample LB-CRC-48-P-01 (TF=0.68) and the normal control LB-Neg-59-P-01 from [35]. For lung cancer, the tumor sample ICD12 (TF=0.53) and the control sample ICT34 were used. For melanoma, the tumor sample ICI06S (TF=0.25) and the control sample ICA079 were used. Area under the ROC curve (AUC) values report the classification performance of Fate-AI and Delfi across the simulated TF, Delta Score is the mean difference between the average prediction scores of tumor and control samples. **B**, Cross-cohort setting: models evaluated on the mixtures were trained on different cohorts with respect to the mixtures. For CRC and lung cancer, models were trained on the Baca et al. cohort [25], for melanoma the model was trained on the Widman et al. cohort [15]. **C**, The Boxplots summarise Delta Scores and ROC AUC of all in silico mixtures (CRC, lung cancer, and melanoma) across different TF levels. **D**, Performance assessment for an in vitro melanoma mixture (n=8) at different TFs tested on models trained on data from this study. C,D: Boxplots show the median as the center, the lower and upper hinges that correspond to the 25th and the 75th percentile, and whiskers that extend to the smallest and largest value no more than 1.5^*^IQR. Values that stray more than 1.5^*^IQR upwards or downwards from the whiskers are considered potential outliers and represented with dots. Significance was computed by a two−sided Wilcoxon rank-sum test (ns: p-value *>* 0.05, ^*^ p-value ≤ 0.05, ^**^ p-value ≤ 0.01, ^***^ p-value ≤ 0.001, ^****^ p-value *<*= 0.0001).

To further validate these findings, we analyzed experimental *in vitro* dilution series. In this setting, pre-treatment melanoma plasma was serially diluted into plasma from healthy donors using plasma-pheresis, generating TF levels between 10^−6^ and 10^−3^. These samples, previously used to validate the MRD-EDGE model [15], which requires matched tumor tissue and ultra-deep WGS (300× coverage), provided an independent benchmark. Fate-AI, trained on data from our cohort of melanoma samples (*n* = 14), accurately distinguished tumor-positive from tumor-negative samples at TF levels as low as 10^−5^ (**Figure 3d**). Collectively, these results demonstrate that Fate-AI, based solely on fragmentomics features, can accurately detect different cancer types with sensitivities ranging from 10^−5^ to 10^−3^.

### Enhanced tumor-naive cancer detection with Fate-AI

We evaluated Fate-AI in various cancer types and different cohorts. Similarly to the simulated case, we examined two scenarios: (i) matched-cohort: models trained and tested on data from the same center, and (ii) cross-cohort: models trained on one cohort and tested on samples from different centers or studies. For benchmarking, we compared Fate-AI with two established fragmentomics-based methods Delfi [18] and Griffin [31]. We also included in the analysis the estimated tumor fraction from IchorCNA [36]. For three cancer types, lung cancer, CRC, and BC, we had samples from two different centers, profiled with both LPWGS and cfMeDIP-seq. Fate-AI, with and without the integration of methylation features derived from the cfMeDIP-seq, achieved the highest area under the curve (AUC) scores (**Figure 4A**). In our cohort, Fate-AI(+Meth) achieved AUCs of 0.89 for BC, 0.88 for CRC, and 0.90 for lung cancer, with sensitivity of 0.88, 0.86, and 0.72, and specificity of 0.89, 0.8, and 0.97 in these three cancer types, respectively. We also tested the models trained on our data on the data from Baca et al. [25], cross-cohort setting. Fate-AI achieved perfect sensitivity and specificity for BC and CRC and an AUC of 0.90 for lung cancer (**Figure 4B**). The better performance in the cross-cohort setting is explained mainly by the higher tumor content of the samples in the test cohort (**Supplementary Figure S1A**). In the cross-cohort setting, we just considered Fate-AI and Delfi. Other methods, including Griffin and IchorCNA, could not be applied because, for the study [25], only the fragment profiles, rather than the raw data, were publicly available, which are required to compute coverage- and copy number–based features. We also generated multimodal data for additional cancer types: melanoma (n=14), PM (n=24), and BLCA (n=38) (**Supplementary Figure S3A**). In the matched-cohort setting, Fate-AI (+Meth) consistently demonstrated an advantage, with an average AUC of 0.91 across the six cancer types (**Figure 4C**). AUC values ranged from 0.84 (BLCA) to 1.0 (melanoma). In comparison, IchorCNA, Griffin, and Delfi exhibited lower performance (mean AUC = 0.70, 0.84, and 0.85, respectively). Fate-AI with methylation features achieved a mean sensitivity of 0.86 and a mean specificity of 0.91 on the six cancer types (**Supplementary Figure S3B**,**C**). Similarly, in the matched-cohort analysis, on the data of [25] including four cancer types: BC (n=15), CRC (n=29), lung cancer (n=38) and PRAD (n=28) versus healthy donors (n=33), Fate-AI(+Meth) achieved an average AUC of 0.97, in contrast, Fate-AI without methylation features had lower discrimination ability of 0.91 (**Figure 4C** and **Supplementary Figure S3B,C,D**). To investigate the relative contribution of cfDNA fragmentomics and methylation features on the observed results, we evaluated the coefficients of the elastic net model. In the fragmentomics-only model, the most influential predictors were measures of differential distribution of fragment lengths and nucleosome-based coverage ratios (**Supplementary Figure S3E**). Specifically, the standard deviation and the total variation consistently showed high weights across multiple datasets, indicating that differences in cfDNA fragmentation patterns, particularly nucleosome positioning and size variability, are major contributors to cancer detection when relying solely on fragmentomics. Upon integrating methylation features, the relative importance of several fragmentation features decreased, while methylation-associated variables emerged as the main predictors in certain cancer classes. Nonetheless, fragment length variability and nucleosome coverage ratios remained among the top features, suggesting complementary information from both data layers. These results demonstrate that cfDNA fragmentomics features alone are highly informative for distinguishing cancer-derived cfDNA, but the addition of methylation signatures enhances classification performance and shifts the balance of predictive importance toward epigenetic features.

**Figure 4.**
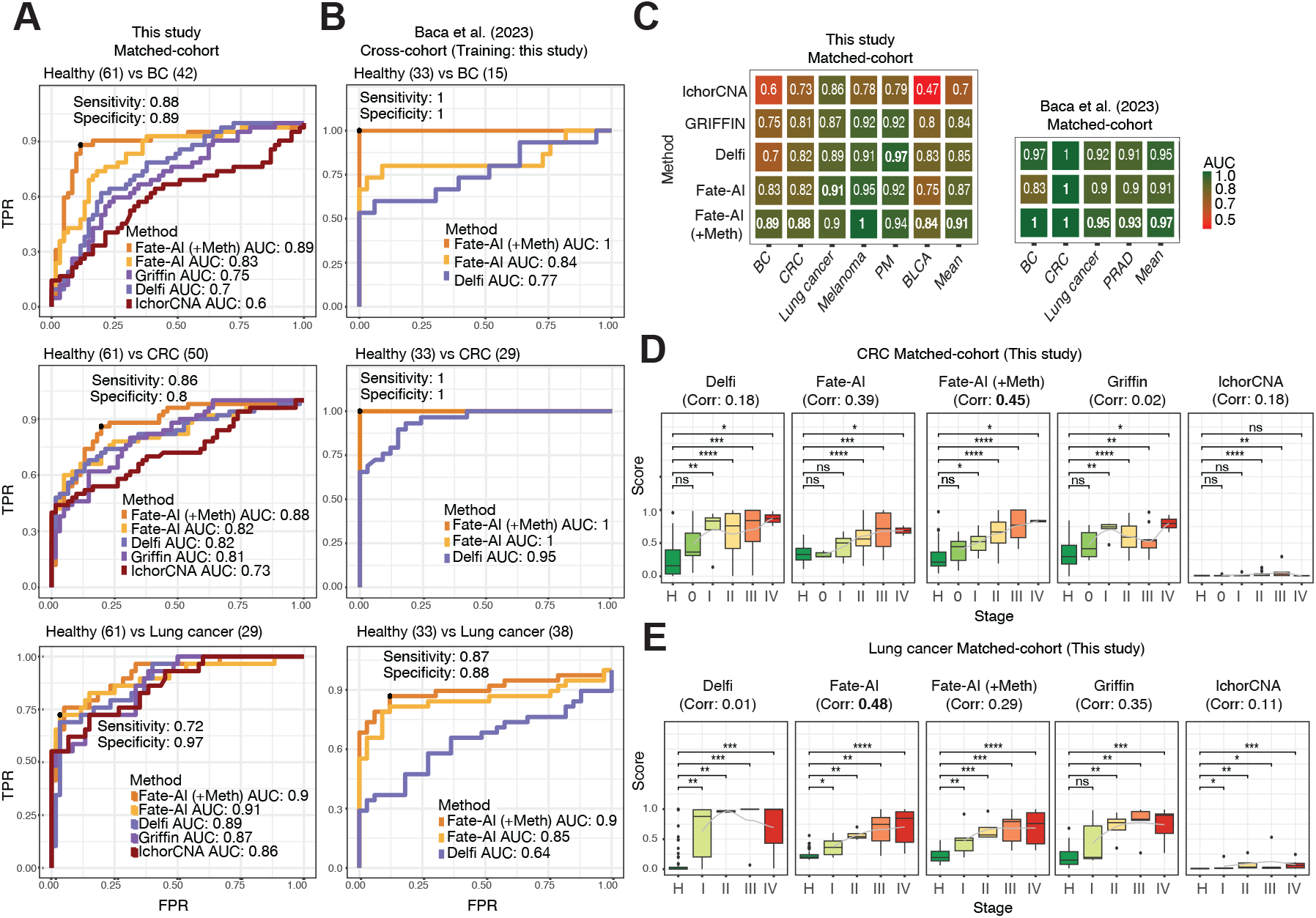
Performance evaluation of methods for cancer detection in WGS-cfMedip. **A**, Matched-cohort setting: models trained and tested on data from this study include 61 healthy donors, 42 BC cases, 50 CRC cases, and 29 lung cancer cases. ROC curves illustrating classification performance (healthy vs. BC, CRC, or lung cancer). The AUC together with the sensitivity and specificity of Fate-AI (+Meth) are indicated. **B**, Cross-cohort setting: models trained on data from this study are tested on the samples from Baca et al. [25] of 33 healthy donors, 15 BC cases, 29 CRC cases, and 38 lung cancer cases. IchorCNA and Griffin could not be evaluated due to the availability of raw data. **C**, Heatmap summarizing the AUC achieved by multiple methods in matched-cohort on this study data (BC n=42, CRC n=29, lung cancer n=29, melanoma n=14, PM n=20, BLCA n=38) and matched-cohort on Baca et al. [25] (BC n=15, CRC n=29, lung cancer n=38, and PRAD n=28). **D**, Boxplots of cancer detection scores stratified by healthy (H) and cancer stage (I–IV) for CRC (n=50), correlation between stage and score of the various models is also reported in the figure title. **E**, same as in D, for lung cancer (n=29). D,E: Boxplots show the median as center, the lower and upper hinges that correspond to the 25th and the 75th percentile, and whiskers that extend to the smallest and largest value no more than 1.5^*^IQR. Values that stray more than 1.5^*^IQR upwards or downwards from the whiskers are considered potential outliers and represented with dots. Significance was computed by a two−sided Wilcoxon rank-sum test (ns: p-value *>* 0.05, ^*^ p-value ≤ 0.05, ^**^ p-value ≤ 0.01, ^***^ p-value ≤ 0.001, ^****^ p-value *<*= 0.0001).

To assess how the model predictions can be used to quantify the amount of disease, we correlated the prediction scores with the tumor stage. In our cohort, both Fate-AI and Fate-AI (+Meth) showed an increase in the prediction score with increasing cancer stage (**Figures 4D,E**). The correlation between the Fate-AI (+Meth) score and cancer stage was 0.45 for CRC (*p*-value = 0.001, Pearson correlation test) and 0.29 for lung cancer (*p*-value = 0.19, Pearson correlation test). For Fate-AI score the correlation was 0.39 for CRC (*p*-value = 0.005, Pearson correlation test) and 0.48 for lung cancer (*p*-value = 0.02, Pearson correlation test). On the other hand, the results from Delfi, Griffin, and IchorCNA were not significantly associated with stage (**Figures 4D,E**): Delfi had a correlation of 0.18 for CRC(*p*-value = 0.22 Pearson correlation test) and of 0.01 for lung cancer (*p*-value = 0.98, Pearson correlation test). Griffin had a correlation of 0.02 for CRC (*p*-value = 0.9, Pearson correlation test) and 0.35 for lung cancer (*p*-value = 0.1, Pearson correlation test). IchorCNA had a correlation of 0.18 for CRC (*p*-value = 0.22) and 0.11 for lung cancer (*p*-value = 0.64, Pearson correlation test). These results show that Fate-AI scores are significantly associated with disease burden.

We also benchmarked Fate-AI on cohorts profiled with only LPWGS: a cohort of 49 patients with PDAC and two public datasets of melanoma [15] and CRC [35]. In melanoma (35 healthy donors vs 38 patients), Fate-AI, trained on our data (cross-cohort setting), achieved an AUC of 0.94, sensitivity of 0.84, and specificity of 0.91, outperforming Delfi (AUC = 0.83) and IchorCNA (AUC = 0.73) (**Supplementary Figure S4A**). Stage-wise analysis further demonstrated the separation of tumor-positive from control samples across disease stages. Fate-AI achieved the highest correlation with stage of 0.44 (*p*-value = 0.006, Pearson correlation test), followed by Delfi with a correlation of 0.25 (*p*-value = 0.09, Pearson correlation test), and IchorCNA with a correlation of 0.2 (*p*-value = 0.24, Pearson correlation test) (**Supplementary Figure S4B**). In the case of PDAC cohort (61 healthy donors vs 49 patients), in a matched-cohort setting, Fate-AI achieved an AUC of 0.97, with a sensitivity of 0.96 and a specificity of 0.90. Similar or slightly lower performance was achieved by Delfi and IchorCNA with an AUC of 0.94 and 0.93, respectively. The correlation with the stage was 0.38 (*p*-value = 6.98*e*^−3^, Pearson correlation test), followed by IchorCNA 0.29 (*p*-value = 0.04, Pearson correlation test) and Delfi 0.17 (*p*-value = 0.23, Pearson correlation test) (**Figure S4C,D**). In the CRC cohort (61 healthy donors vs 43 patients) of Hallermayr et al. [35], cross-cohort setting, Fate-AI achieved an AUC of 0.76, with a sensitivity of 0.81 and a specificity of 0.62 (**Supplementary Figure S4E,F)**. IchorCNA achieved slightly better performance with an AUC of 0.81. Fate-AI achieved the highest correlation with stage of 0.61 (*p*-value = 1.38*e*^−5^, Pearson correlation test), followed by Griffin 0.58 (*p*-value = 4.74*e*^−5^, Pearson correlation test), IchorCNA 0.53 (*p*-value = 0.0003, Pearson correlation test), and Delfi 0.55 (*p*-value = 0.0001, Pearson correlation test).

Collectively, these findings highlight the sensitivity, robustness, and generalization of Fate-AI models, particularly at low tumor fractions and in early-stage disease, both with LPWGS features alone and when incorporating cfMeDIP features.

### Fate-AI allows accurate longitudinal disease progression monitoring

Beyond detection at baseline, a crucial test of any cfDNA-based model is its ability to monitor patients longitudinally, capturing changes in tumor burden during treatment and follow-up. Hence, we next assessed the ability of Fate-AI to capture disease dynamics over time in colorectal cancer, where recurrence often arises from minimal residual disease that remains undetectable by conventional imaging or biomarkers. Therefore, longitudinal monitoring of circulating tumor DNA is crucial for capturing early molecular relapse and guiding therapeutic intervention. We applied Fate-AI, trained on our internal CRC, to 198 longitudinal samples from 50 patients treated with chemotherapy [35] (1-23 samples per patient, average: 4). Because only LPWGS data were available, Fate-AI with fragmentomics-derived features was used. We tracked scores from four distinct methods, Fate-AI, Delfi, Griffin (trained on our cohort), and IchorCNA, over time. The output of the four methods was cross-referenced with treatment events, such as surgery and chemotherapy initiation, as well as disease status categories, including remission, disease progression, and death. Two illustrative cases are reported in **Figures 5A,B**. For patient LB-CRC-32, Fate-AI exhibited dynamic changes corresponding to clinical transitions, particularly aligning with partial remission and disease progression events (**Figure 5A**). Qualitatively, the Delfi and Griffin scores showed high variability over time, which did not always correspond to the response to treatments. IchorCNA showed a generally flat prediction, although low in almost all time points, without an increase in clinical responses related to disease progression. A similar trend was observed for the patient LB-CRC-34, where Fate-AI scores followed the patient’s clinical trajectory (**Figure 5B**). The Fate-AI score decreased significantly near the partial remission stage, indicating a reduction in tumor-derived signals in circulation. After the first interruption of chemotherapy, the score increases, aligning with the expanded residual disease. Following the second chemotherapy cycle, the score dropped again, suggesting a positive treatment response. However, at disease progression, the Fate-AI score sharply increased. IchorCNA showed a good correlation with staging and therapy-related information, even though scores remained relatively low. On the contrary, the dynamic changes of Delfi and Griffin seemed to follow the disease trajectory to a lesser extent. Additional cases, with similar overall behavior, are reported in **Supplementary Figure S5**.

**Figure 5.**
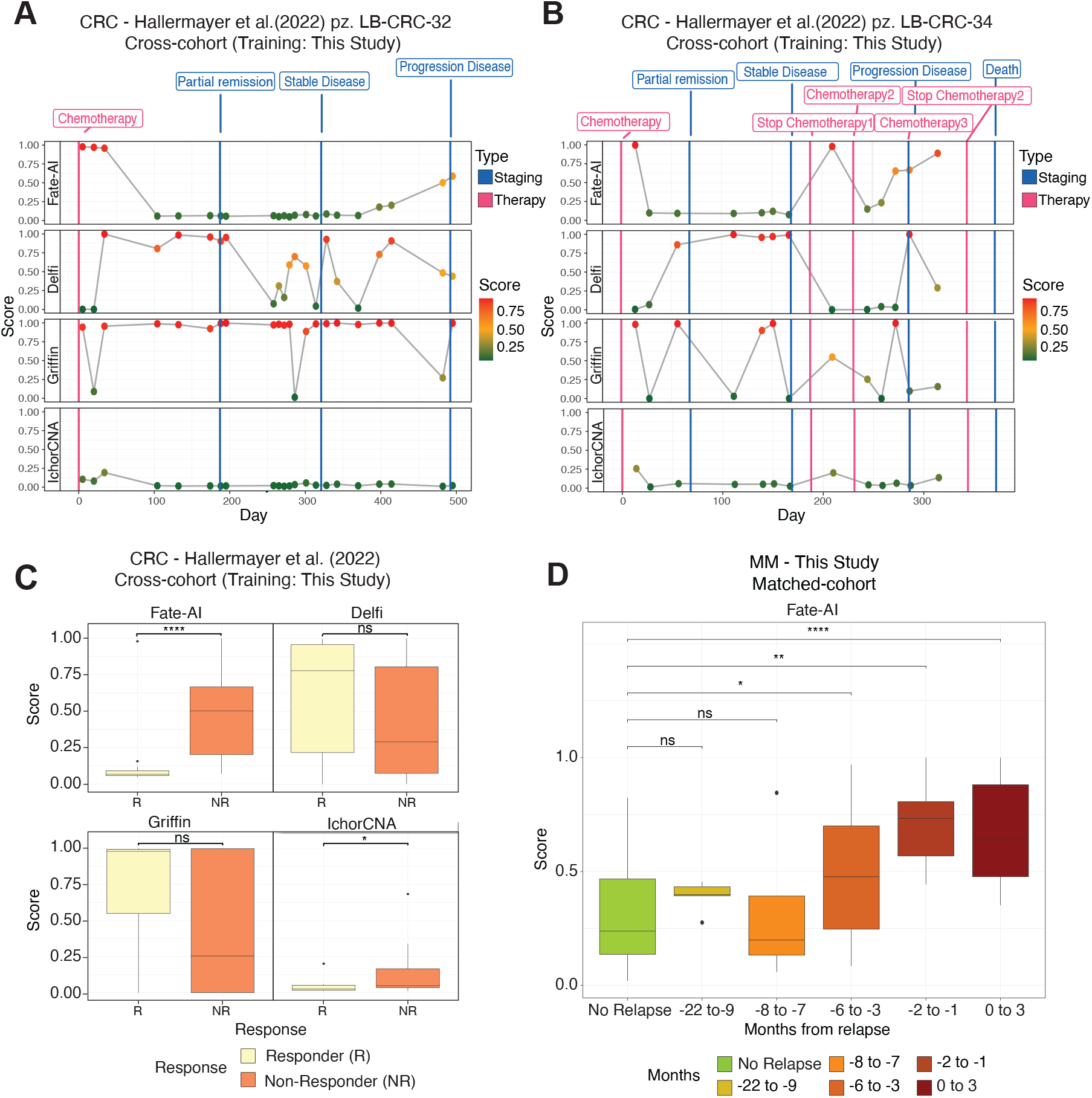
Longitudinal disease progression monitoring. **A**, The model trained with data from this study was applied to 198 longitudinal samples from 50 patients (average four samples per patient, range 1-23) treated with chemotherapy [35]. The *x* axis represents the time, starting at the first treatment, and the *y* axis reports the score of Fate-AI (fragmentomics features only), Delfi, Griffin, and IchorCNA for patient LB-CRC-32. Treatments are reported as red lines, and staging events as blue lines. **B**, Same as in A, for patient LB-CRC-34. Additional cases are reported in the supplementary figure S5. **C**, Scores from multiple methods (Fate-AI, Delfi, Griffin, IchorCNA) for post-treatment samples of the CRC cohort [35] stratified by response. **D**, Boxplots showing Fate-AI scores in post-treatment patients with MM stratified (n=91) by time to relapse (No relapse, −22 to −9 months, −8 to −7 months, −6 to −3 months, −2 to −1 months, and 0 to 3 months). C,D: Significance was computed by a two−sided Wilcoxon rank-sum test (ns: p-value *>* 0.05, ^*^ p-value ≤ 0.05, ^**^ p-value ≤ 0.01, ^***^ p-value ≤ 0.001, ^****^ p-value *<*= 0.0001). Boxplots show the median as center, the lower and upper hinges that correspond to the 25th and the 75th percentile, and whiskers that extend to the smallest and largest value no more than 1.5^*^IQR. Values that stray more than 1.5^*^IQR upwards or downwards from the whiskers are considered potential outliers and represented with dots.

To enable a quantitative comparison of disease scores across the evaluated methods, the 198 plasma samples were classified according to the nearest clinical assessment (within *±*50 days). Cases corresponding to stable disease (SD), partial response (PR), or complete response (CR), contributing to the disease control rate, were classified as responders. Those with progressive disease (PD) were classified as non-responders. Fate-AI demonstrated significant discrimination between responder (R) and non-responder (NR) groups. (*p*-value = 5.7 × 10^−6^, Wilcoxon rank-sum test ; **Figure 5C**), a performance that was also observed, though to a lesser extent, with IchorCNA (*p*-value = 0.01, Wilcoxon rank-sum test).

We also evaluated Fate-AI on liquid cancers, with a particular focus on MM. Relapse in MM is almost inevitable, with most patients eventually developing aggressive and treatment-refractory disease [37]. Current clinical monitoring relies primarily on invasive bone marrow biopsies and serum biomarkers, which often fail to capture the full range of clonal heterogeneity or to provide timely detection of emerging resistant clones. Therefore, Fate-AI may provide a powerful tool for early relapse detection, real-time disease monitoring, and improved risk stratification in MM. We profiled a cohort of 24 MM patients enrolled in a clinical trial investigating maintenance therapy with lenalidomide after induction therapy (NCT02538198) [38, 39, 40]. Ninety-one samples were collected at up to four timepoints (with an average of 3.8 samples per patient, a minimum of 2, and a maximum of 4), before, during, and after maintenance therapy, at a median interval of 167 days (range: 12–616 days). We compared patients who did not experience progression (12) with those who did progress (12) after maintenance therapy. Longitudinal patient trajectories show that Fate-AI scores remain low in non-progressors but increase months before clinically documented progression (**Supplementary Figure S6**). Consistently, aggregated analyses revealed significant differences as early as 6 months prior to relapse (**Figure 5D**). Finally, we also performed a longitudinal analysis in post-treatment samples of pediatric tumors. We considered a cohort of patients with ES [21]. Available data include 66 samples at diagnosis, 82 samples under treatment, 17 samples after relapse, and 14 samples after remission from 84 patients (with an average of two samples per patient, ranging from 1 to 10). Fate-AI scores, for post-therapy samples, demonstrated distinct trajectories between patients who later relapsed and those who remained in remission (**Supplementary Figure S7A**). Starting from months 1-3, scores began to diverge, with relapse patients maintaining consistently high values while remission patients showed a progressive decline (*p*-value = 0.070, Wilcoxon rank-sum test). This separation became statistically significant at months 4–6 (*p*-value = 0.018, Wilcoxon rank-sum test), highlighting a predictive window for the detection of relapse. Beyond 6 months, patients who relapsed sustained elevated scores (0.9–0.95), whereas remission patients remained at lower levels. Interestingly, when considering the pre-therapy samples of the same cohort, Fate-AI score showed a significant difference between patients who relapsed (*p*-value = 0.012, Wilcoxon rank-sum test, **Supplementary Figure S7B**). This difference, instead, was not evident in the samples just after the start of the therapy as shown in the first points of the curves in the Supplementary Figure S7A.

Together, these analyses establish Fate-AI as a robust cfDNA framework capable of tracking tumor burden across both solid and hematological malignancies. In MM, Fate-AI predicted relapse months before clinical progression, while in CRC and ES, it dynamically reflected treatment response and significantly distinguished R from NR.

### Using the Fate-AI score to predict response to immune therapy

We then asked if the Fate-AI score could be used in specific therapeutic contexts, such as immune therapy, where the response depends both on tumor-intrinsic factors (tumor mutational burden, neoantigen load, MSI status, and others), characteristics of the microenvironment, such as the presence of adaptive antitumor immunity and cytotoxic lymphocytes, and other environmental factors. We applied Fate-AI in a cohort of 20 cases before treatment of PM treated with immunotherapy in the second line [41], Fate-AI (+Meth) disease scores were significantly different between patients who responded versus patients who did not respond (*p*-value = 0.02, Wilcoxon rank sum test) (**Supplementary Figure S7C**). This result is in line with the recent finding that epigenetic subtypes are associated with response to immunotherapies in mesothelioma [41]. We also considered a set of 38 pre-treatment and 95 post-treatment plasma samples from 40 melanoma patients (average three samples per patient, minimum 2, maximum 5) reported in [15]. Our Fate-AI melanoma model, trained on samples from this study and [25], cross-cohort setting, was then tested on the post-treatment cases. Fate-AI scores were significantly associated with patient outcomes (**Supplementary Figure S7D** left panel). Specifically, patients who achieved CR or PR showed significantly lower scores as compared with those with PD. Similarly, patients who experienced a progression after immunotherapy displayed significantly higher Fate-AI scores relative to those who remained progression-free (*p*-value = 0.038, Wilcoxon rank-sum test) (**Supplementary Figure S7D** right panel). Overall, these observations suggest that the Fate-AI score may capture biological characteristics relevant to the response to immune therapy.

### Tissue-of-origin identification with Fate-AI

Features such as fragment length distributions, end motifs, nucleosome positioning patterns, and coverage profiles across the genome reflect the epigenetic and chromatin organization of the tissue from which the DNA originated [16]. Since the Fate-AI uses tumor-specific features from the Fragment Differential Distribution, we evaluated its ability to classify the tissue of origin (TOO) of tumors. By design, the FDD has different shapes for different cancer types; in particular, it presents sharper transitions for the cancer of interest, facilitating the TOO discrimination (**Figure 6A**). We focused on six tumor types (BC, CRC, ES, lung cancer, Melanoma, and PDAC), each represented by at least ten samples with a TF greater than 0.10 across the available cohorts. To train the model, we just concatenated the tissue-specific features (Methods). Multiclass Receiver Operating Characteristic (ROC) curves demonstrated area under the curve (AUC) values ranging from 0.84 (BC) to 0.97 (CRC) (**Figure 6B**). CRC and Melanoma samples displayed highly distinctive CNA-associated fragmentation profiles, whereas other tumor types exhibited partial overlap (**Figure 6C**). These results suggest that while certain tumors harbor unique CNA-driven fragmentomic patterns that facilitate classification, others share broader genomic alterations that may reduce specificity.

**Figure 6.**
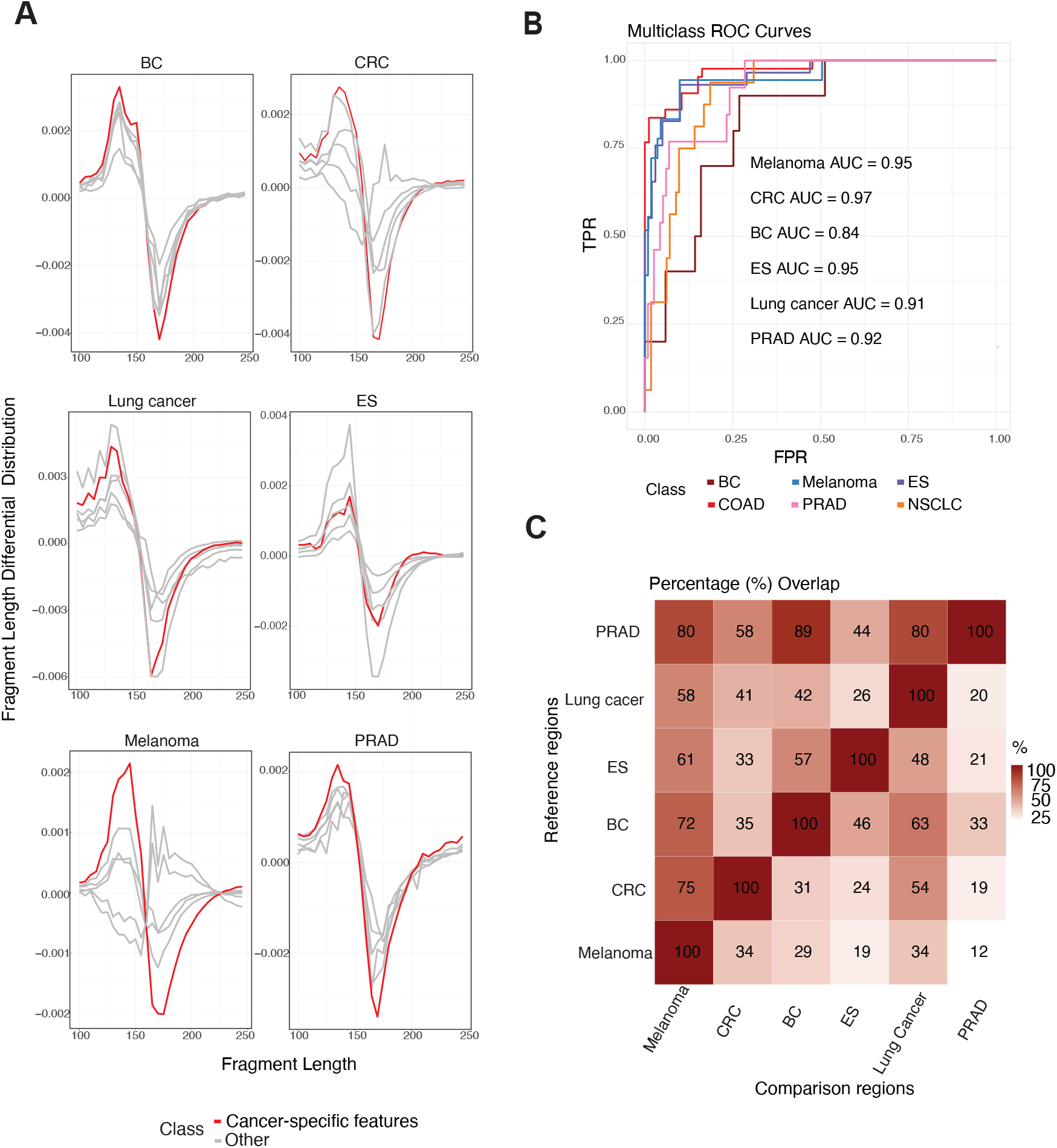
Tissue-of-origin classification performance. **A**, Fragment Length Differential Distribution for each cancer type across datasets stratified by CNA-specific tumor type. Red curves correspond to median FDD from samples of the tumor type whose CNA regions were analyzed, while gray curves represent other tumor types. **B**, Multiclass ROC curves illustrating classification performance of tissue-of-origin, for six tumor types with at least ten samples having TF *>* 0.10 (BC n=11, CRC n=45, ES n=29, lung cancer n=16, melanoma n=19, and PRAD n=13), from this study and external datasets ([25, 35, 21, 15]). **C**, Heatmap of pairwise percentage overlap between CNA regions [34] across different tumor types. The color scale represents the degree of overlap, ranging from low (white) to high (red), with values shown inside each cell. Reference regions (rows) are compared against Comparison regions (columns). Values indicate the fraction of Reference regions overlapping with each Comparison set.

## Discussion

Liquid biopsy has become a central weapon in precision oncology, offering non-invasive access to tumor-derived genetic and epigenetic information through analysis of cell-free DNA (cfDNA) [42]. Multiple classes of assays have been developed, each with strengths and limitations. Despite its potential, the clinical deployment of cfDNA-based tumor-agnostic assays has been hampered by variability in sequencing protocols, batch effects, and the limited generalizability of existing models across heterogeneous cohorts. Current strategies based on mutations, copy number alterations, or fragmentomic features have achieved proof-of-concept success, but their robustness often diminishes when applied across datasets or at low tumor fractions, particularly in early-stage disease [18, 15] Tumor-informed mutation panels, relying on bespoke sequencing of patient-specific variants, achieve high specificity but are constrained by sampling depth, tumor heterogeneity, and acquired resistance. Tumor-naive fragmentomic methods [18], end-motif profiling [19], and nucleosome positioning analysis [20] have been proposed as innovative strategies for non-invasive cancer detection and disease measurement. Similarly, methylation profiling has shown success in identifying renal [22], prostate [23], and intracranial tumors [25]. Fate-AI combines these orthogonal signals into a unified knowledge-informed model, coupled with a feature selection based on contrasts on the same sample, to ensure robust generalization. This strategy aims to combine the advantages of tumor-naive approaches with the specificity of tumor-informed approaches, while minimizing their drawbacks. To benchmark Fate-AI, we generated multiomic profiles of 432 plasma samples from eight cancer types (BC, CRC, MM, lung cancer, Melanoma, PM, BLCA, and PDAC). We validated its generalization performance across multiple independent cohorts encompassing 787 samples. We achieved tumor detection performance up to tumor fractions of 10^−5^ in experimental dilution series and 10^−3^ for in silico admixtures. This level of sensitivity is particularly relevant for minimal residual disease monitoring, where ctDNA burden is often below the detection threshold of mutation-based assays or when tumor-informed panels fail due to tumor evolution and acquired resistance.

The correlation between Fate-AI scores and disease stage, as well as their dynamic modulation during therapy, supports the use of Fate-AI as a quantitative biomarker of tumor burden. Its ability to detect molecular relapse months before clinical progression underscores its potential for minimal residual disease (MRD) surveillance and real-time treatment monitoring, particularly in settings where imaging or protein biomarkers lack sensitivity.

Fate-AI enabled accurate classification of tumor tissue of origin, achieving AUC values between 0.84 and 0.97 across different cancer types. While some tumor types (e.g., melanoma and CRC) displayed highly distinctive profiles, others exhibited partial overlap, reflecting the shared genomic and epigenomic architecture across malignancies.

Nonetheless, multiple limitations remain. While the region-based normalization mitigates technical variability, tumors with low ctDNA yields or rare CNA/methylation events may remain challenging. Furthermore, prospective clinical trials with larger, more diverse cohorts are necessary to validate Fate-AI’s performance in real-world settings and to assess its integration into clinical decision-making.

In conclusion, Fate-AI, by adopting a “knowledge-informed” approach, tries to overcome the limitations of both the tumor-informed strategies, which need original tissue, and tumor-agnostic approaches that can lack generalizability. Its performance across tumor types, stages, and datasets evidences its translational potential for early detection, MRD monitoring, and tissue of origin classification. These findings support the further development of multimodal cfDNA frameworks as clinically actionable tools in precision oncology.

## Methods

### Sample collection

This study was approved by the relevant local ethics committees and institutional review boards and was conducted in accordance with the Declaration of Helsinki protocol. Samples were collected from different centres as listed in **Supplementary Table 1**. 50 CRC and 42 BC patients were recruited at Istituto Oncologico Del Mediterraneo (IOM), and the studies were approved by the Ethics Committee of Catania 2 with (protocol n:436/C.E. and n:117/C.E). 29 NSLC patients were enrolled at Azienda ospedaliera universitaria (Vanvitelli), Italy. 38 BLCA patients were enrolled at Azienda ospedaliera universitaria (Vanvitelli) and Azienda Ospedaliera dei Colli Monaldi, Italy. The study was approved by the Ethical Committee of Azienda Universitaria Policlinico of the University of Campania “L. Vanvitelli”(protocol n. 6691). 49 PDAC patients were enrolled at Miller School of Medicine, University of Miami, USA (protocol n.20230674). 40 Melanoma samples derived from the phase Ib NIBIT-M4 study (NCT02608437) in accordance with the ethical principles of the Declaration of Helsinki and the International Conference on Harmonization of Good Clinical Practice. For available matched melanoma cases, we collected samples after 4 weeks and 12 weeks from therapy (Ipimilumab +Guatecitabine). 20 PM patients were recruited from the NIBIT-MESO-1 (NCT02588131). 91 MM patients were enrolled at Memorial Sloan Kettering Cancer Center(protocol n.15-129 A(22)), and 69 healthy individuals without cancer were enrolled from these centers as controls. Peripheral blood was collected in EDTA-containing tubes and plasma was separated within 1.5 hours of collection. Plasma was aliquoted and stored at -80 °C until processing.

### cfDNA isolation

MM plasma samples from Memorial Sloan Kettering Cancer were thawed in a 37°C dry bath for 30 minutes, cfDNA isolation was performed using QIAamp MinElute ccfDNA Midi Kit (Qiagen) on the QIAcube instrument, and DNA was eluted in 20 *µl* nuclease-free water. For all the others samples, cfDNA isolation was performed using Qiamp Circulating acid kit (Qiagen) and elute in 50*µl* of AE buffer. Double-stranded DNA was quantified using the Qubit dsDNA HS Assay (Invitrogen), and cfDNA was stored at -80 °C.

### LP-WGS libraries preparation

Libraries preparation for MM cohort was performed using NEBNext UltraTM II DNA,while for the others cancer types, the Kapa Hyper prep kit (Roche) was used following the manufactor’s instructions. Libraries were amplified using 8-10 cycles and purified with AMPure XP beads (Beckman Coulter). Final libraries concentration was examined using the Qubit dsDNA HS Assay (Invitrogen), and fragment size quality check was visualized on 2100 Bioanalyzer Biosystem (Agilent).

### cfMeDIP-seq

cfMEDIP-seq protocol was performed as described previously[43]. Briefly, CfDNA libraries were proceed using the Kapa Hyper Prep kit (Roche) and adaptors from NEB next multiplex oligos for Illumina kit (NEB) were added. Lamba DNA, a mix of unmethylated and methylated PCR amplicons, was added to uncomplete libraries. Methylated and unmethylated Arabidopsis thaliana DNA was added for quality control purposes (Diagenode). One part of the libraries was kept as input control (IC) and while the rest of the libraries were immunoprecipitated with antibody 5mC (33Dclone) (IP) using MagMeDIP-kit (Qiagen). The efficiency of the immunoprecipitation was verified via qPCR by detecting the recovery of the spiked-in Arabidopsis thaliana DNA (both methylated and unmethylated), following Diagenode’s instructions. Libraries were amplified using between 9-15 cycles with the following protocol: 98°C for 3min, 98°C for 20s, anneal at 65°C for 15s, and extension at 72°C for 30s. Complete libraries were purified with AMPure XP beads (Beckman Coulter). Quality was examined by capillary electrophoresis utilizing a 2100 Bioanalyzer Biosystem (Agilent).

### Sequencing

LPWGS and cfMeDIP-seq libraries were independently pooled based on expected final coverage and sequenced across multiple flow cell lanes to minimize lane-to-lane variability in yield. Sequencing was performed on an Illumina NovaSeq 6000 platform using a paired-end 2 × 150 bp read configuration. For LPWGS data, the average coverage (mean *±* SD; range) across cancer types and healthy samples was as follows: BC, 4.89 *±* 1.71× (2.25–8.94); CRC, 4.81 *±* 1.55× (1.31–7.44); lung cancer, 2.49 *±* 0.90× (0.90– 4.76); melanoma, 8.23 *±*2.58× (4.65–19.2); PM, 8.14 *±*1.12× (6.19–11.1); MM, 5.52 *±*2.16×(0.72–13.3); PDAC, 19.9 *±* 3.14× (14.9–26.4); BLCA, 5.17 *±* 0.98× (2.29–6.80); and healthy controls, 4.74 *±* 3.09× (0.54–17.3). For cfMeDIP-seq data, the average number of reads per sample (mean *±* SD; range, in millions) across cancer types and healthy samples was as follows: BC, 45.3 *±* 19.2 M (15.6–93.1); CRC, 50.1 *±* 20.8 M (14.2–94.4); lung cancer, 31.0 *±* 16.5 M (7.34–73.2); melanoma, 39.2 *±* 27.9 M (6.96– 103.0); PM, 31.7 *±* 18.6 M (3.4–71.6); BLCA, 70.1 *±* 37.4 M (11.1–149.0); and healthy samples, 73.7 *±* 44.7 M (11.2–161.0).

### Fragmentomics Features

The sequencing reads obtained from the Low pass whole genome sequencing (LPWGS) of liquid biopsy were aligned with respect to the human genome hg38 using Sentieon bwa-mem (0.7.17-r1188), then duplicate reads were removed and finally, the indel realignment and base recalibration were performed using Sentieon tools v. 202112.

As a preprocessing step we apply GCparagon (v0.5.4) to correct GC content and fragment length biases by quantifying their effects on read coverage and applying length-specific normalization and statistical corrections, resulting in a bias-corrected profile [44].

Then, genomic regions with known copy number alterations are extracted from the Progenetix Database [34], for each cancer type. Recurrence thresholds were selected according to the percentage of covered samples as a function of the size of the selected genomic regions as depicted in Figure 1C and listed in Supplementary Table S3. We categorized the expected Copy Number Variation (CNV) regions between ctDNA and cfDNA into two main groups: expected Gain Copy Number Regions (eGCR), where a high presence of ctDNA (high TF) is anticipated, and expected Loss Copy Number Regions (eLCR), where, in contrast, a low presence of ctDNA (low TF) is expected.

The eGCR and eLCR regions were partitioned into non-overlapping bins of 3,000,000 bp. Within each bin, we quantified fragment and coverage-based metrics, including mean fragment length, coverage, nucleosome core coverage (140–159 bp), chromatosome coverage (160–170 bp), nucleosome coverage (171–240 bp), the ratio of nucleosome core to nucleosome coverage, and the ratio of (nucleosome core + chromatosome) to nucleosome coverage. For each metric *m*, the empirical cumulative distribution function (ECDF), 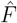, was computed separately across bins in the eGCR and eLCR regions. TheFragment Differential Distribution (*FDD*) was then defined as:

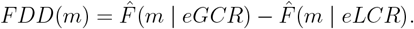

Additionally, we compute the *FDD* for global fragment lengths without binning. We then use as features the total variation of *FDD, TV* (*m*) = Σ |*FDD*(*m*)|, and its standard deviation 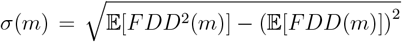 for each global and local metric. Furthermore, for global fragment lengths, we also compute the Kullback–Leibler divergence (KL) between the two distributions underlying the *FDD*.

### Methylation features

The contrast in methylation profiles between tumor-derived and plasma-derived cfDNA allows for the identification of tumor-specific signals and to compare it against the cfDNA background of healthy cells. This is achieved by focusing on genomic regions that can significantly differ between tumor and normal tissues, and on those known to be hypermethylated, typically in the healthy cells we find in plasma. Since cfMeDIP-seq captures only methylated cfDNA fragments, we specifically focus on hypermethylated regions for analysis, excluding hypomethylated regions from consideration. This ensures that our approach directly leverages the signal present in the data, allowing for a more accurate and biologically meaningful interpretation of the methylation landscape. We categorized the expected differentially methylated regions (DMRs) between ctDNA and cfDNA, respectively, into two main groups: expected Hypermethylated Regions in Tumor (eHRT) and expected Hypermethylated Regions in Plasma (eHRP). The first set, eHRT, was obtained through an analysis of DMRs between primary tumors and normal solid tissues, which was carried out for each cancer type using the Cancer Genome Atlas (TCGA) [45]. From the most significant regions (Fold-change *>* 1.5 and adjusted *p*-value *<* 0.01, Wilcoxon rank-sum test with Benjamini-Hochberg correction), we selected up to 3,000 DMRs as those expected to be hypermethylated in cancer. Regions Hypermethylated in Plasma are instead typically associated with hypermethylated in cell types found in plasma, including vascular endothelial cells, hepatocytes, erythrocyte progenitors, monocytes, neutrophils, B cells, CD4+ T cells, CD8+ T cells, and NK cells [46]. The second set, eHRP, thus was obtained using Methylation Atlas Deconvolution [46]. DMRs between plasma-associated cell types and other cell types in the dataset were obtained as explained above and, similarly, up to 3,000 DMRs as those in the set of eHRP. For both sets, DMRs were defined as regions of 300 bp centered on the CpG array-based probe coordinates in the TCGA and Methylation Atlas datasets. We then compute three features for each cfMeDIP-seq profile, summarizing the observed methylation signal in the regions enriched in ctDNA and those depleted in ctDNA:

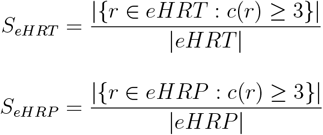

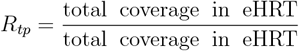

where *c*(*r*) is the read count in region *r*. These features represent the percentage of regions in the **sample with at least three reads in eHRT, eHRP, and the ratio between the overall coverage of these** two sets of regions.

### Predictive Models

The feature matrix, based on fragmentomics features or combined with methylation features, was used as input for a logistic regression-based classifier to discriminate between healthy controls and cancer patients. Specifically, we used an elastic net model implemented in the glmnet package, with *ε* = 0.5 to balance L1 (Lasso) and L2 (Ridge) regularization. The *λ* parameter, which controls the strength of regularization, was explored over a range from 0 to 0.05 in increments of 0.01. A 10-fold cross-validation was used to fine-tune model parameters. For matched-cohort scenarios, model performance was assessed within the same cohort using 100 repetitions of 10-fold cross-validation to ensure robustness and reliability, and to prevent performance fluctuations due to data partitioning. The final performance estimates were obtained by averaging prediction results across all repetitions [18]. For cross-cohort scenarios, an independent cohort of the same cancer type was used to train the logistic regression–based classifier, while the cohort under assessment served as the test set for score prediction. When our dataset was used for training, we used tumor samples with a tumor fraction (TF) greater than 0.03 and healthy control samples with a TF below 0.01.

### In silico TF admixtures

To assess cancer detection we generate in silico admixtures at different level of Tumor Fraction using whole-genome sequencing (WGS) data from plasma samples across multiple cancer types (lung cancer, CRC, Melanoma) with high-TF and healthy samples. The approach ensures unbiased mixing by pairing cancer samples with non-cancer plasma samples from the same sequencing center and platform [15].

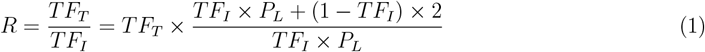

Where *TF*_*I*_ represents the initial Tumor Fraction in the high-TF cfDNA cancer sample, *TF*_*T*_ is the target Tumor Fraction, and *P*_*L*_ denotes the ploidy of the tumor sample. We obtain the fraction of reads to be extracted from each sample of the pair in the following way:

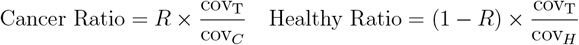

where cov_*C*_ and cov_*H*_ are the read depths of cancer and healthy samples, respectively, and cov_T_ is the targeted sequencing depth for the final in silico admixture sample. These formulas ensure proper dilution of high-burden cfDNA and normalization of sequencing depth across different tumor fractions. SAMtools (v. 1.19.2) was used to downsample (view -s) and admix (merge) cancer plasma and healthy control plasma cfDNA reads accounting for estimate Ratio, to obtain the Target Tumor Fraction (*TF*_*T*_ ). To improve the statistical reliability of the benchmark on detection, 10 independent replicates of the in silico admixture are generated, alongside multiple in silico replicates of the healthy control sample. We generate independent replicates at 10 different levels of targeted TF: 10^−6^, 10^−5^, 10^−4^, 10^−3^, 5 × 10^−3^, 10^−2^, 2×10^−2^, 3×10^−2^, 5×10^−2^ and 10^−1^. Replicates of the healthy sample are obtained by subsampling with a sequencing coverage of 90%. This approach helps establish accurate detection thresholds for circulating tumor DNA (ctDNA) across different cancer types by minimizing background noise and ensuring robust analytical performance.

### MRD Detection and Longitudinal Dynamics

For the evaluation of minimal residual disease, we applied Fate-AI together with other methods (Delfi, Griffin, IchorCNA) to the longitudinal plasma sample of CRC reported in [35], using the cross-cohort setting (“Predictive Models”). Each model was trained on data from this study. For each patient in the test cohort, all available time points were considered and cross-referenced to clinical metadata, including treatment events (surgery, chemotherapy, immunotherapy) and disease status categories (remission, PR, progression, death). The MRD scores dynamics of each method were aligned with the treatment events and disease outcomes to assess the temporal concordance between the predicted tumor burden and clinical course. For quantitative assessment in the MRD context, plasma samples were labeled with respect to the closest disease status using a maximum window of range ±50 days. Plasma samples labeled as remission, PR, or SD were grouped as responders, while those labeled as PD were grouped as non-responders.

Fate-AI was further assessed on longitudinal multiple cancer cohorts. In the cross-cohort setting (train data from this study and Baca et al. [25]) on conventional immunotherapy melanoma samples from Widman et al. [15], where the statistical comparison between responder categories and between plasma samples of patients with disease progression or death event (PFS) or not was performed.

In the matched-cohort setting for MM, where no pre-treatment samples were available, the positive class (cancer) of the training was defined as the set of plasma samples with at least one positive test result (MRD+, M-Protein, Light Chain, or PET). In contrast, we used as control the plasma samples with TF *<* 0.02 and no evidence of future relapse.

In the MRD analysis of PM patients from this study, the matched-cohort setting was used to train the model with cross-validation.

For ES from Pender et al. [21], the matched-cohort setting was employed, using pre-treatment samples with control samples to train the model and post-treatment samples as the test set. Statistical comparisons between different categories of response to treatment or future progression/relapse were performed using a two-tailed Wilcoxon test.

### Tissue-of-origin Identification

To train Fate-AI tissue of origin classification from plasma samples, tissue-specific features were derived for each cancer type based on region-specific copy number alterations (CNAs) obtained from the Progenetix Database [34], as before. For this assessment, the union of these tissue-specific feature sets across all cancer types was then concatenated into a single feature matrix. A multinomial logistic regression-based classifier as described above (“Predictive Models”). The model’s performance was evaluated using 100 repetitions of 10−fold cross-validation iterations to ensure robustness and reliability. To assess the discriminative performance in each cancer type, Receiver Operating Characteristic (ROC) analysis was conducted in a one-vs-all framework using the pROC package.

## Data and code availability

Raw data (cfMeDIP-seq and LPWGS) generated for this study will be deposited on EGA before acceptance. Public raw data (WGS) are available on EGA: Melanoma (EGAD00001010926)[15], healthy and in vitro mixture of Melanoma (EGAD00001011352)[15], ES (EGAD00001007080)[21], CRC (EGAD00001009309)[35]. BED files containing genomic alignments (cfMeDIP-seq and LPWGS) are available on GEO (GSE243474)[25].

Details and codes for the data processing and annotation is available on GitHub (https://github.com/ceccare AI).

## Acknowledgements

Research reported in this publication was supported by the Biospecimen Shared Resource (BSSR, RRID: SCR 022889), the Oncogenomics Shared Resource (OGSR), and the Biostatistics and Bioinformatics Shared Resource (BBSR, RRID: SCR 022890) of the Sylvester Comprehensive Cancer Center at the University of Miami, RRID: SCR 022502, which is supported by the National Cancer Institute (NCI) of the National Institutes of Health (NIH) under award number P30CA240139.

## Supplementary Figure Legends

**Figure S1.**
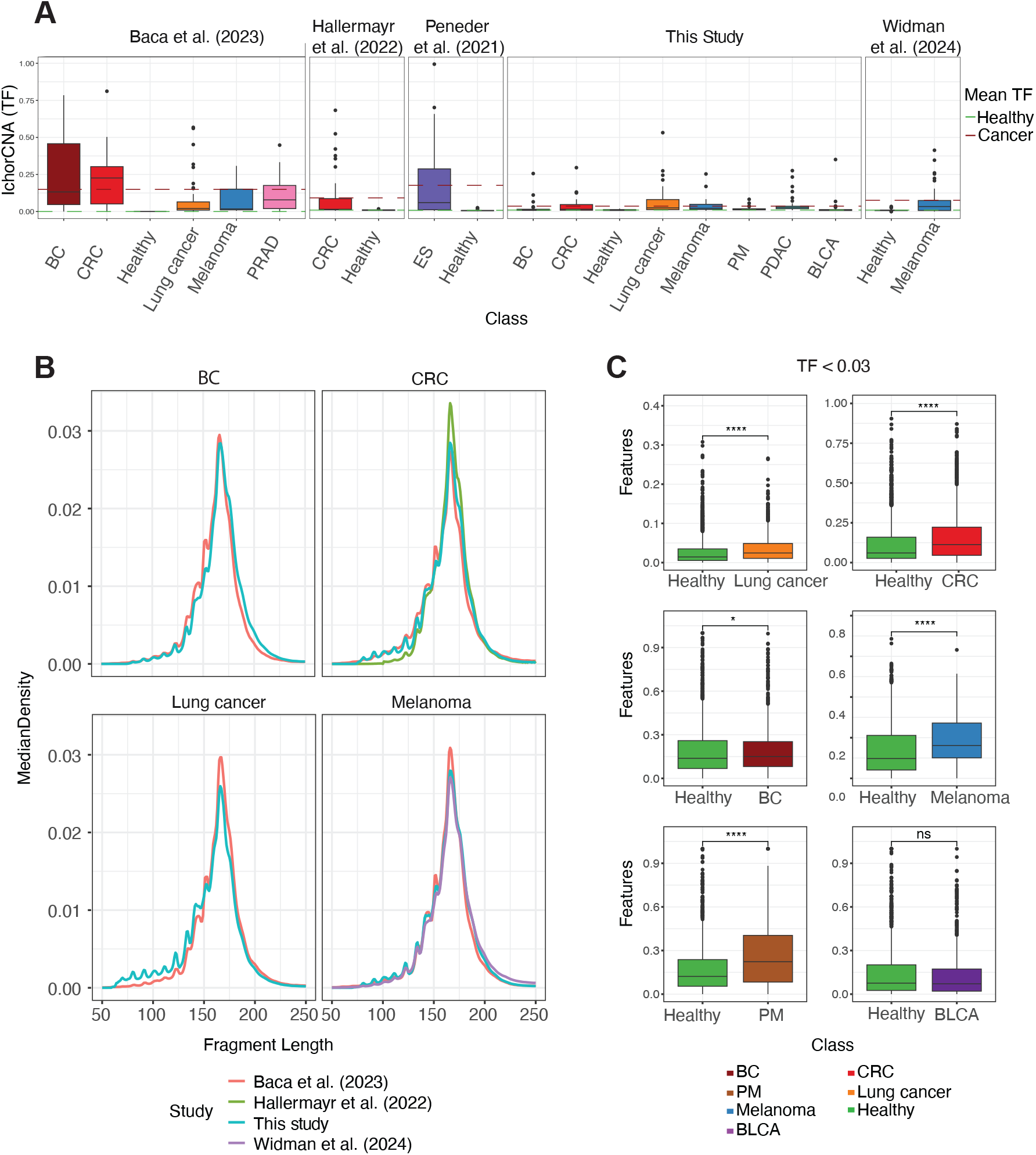
Technical variations effect on Fragment Length Distribution and the distribution of features in early-stage. **A**, Boxplots show the distribution of the tumor fraction (TF) estimated by IchorCNA[36] across multiple studies. The dotted lines show the mean tumor fraction of healthy samples (green) and tumor samples (red) in each study. **B**, Density of cfDNA fragment lengths observed in cancer samples across multiple studies. **C**, Boxplots show the distribution of Fate-AI features (Methods) in healthy controls and early-stage cancer samples with low TF less than 0.03. Statistical comparisons were performed using the Wilcoxon rank-sum test (ns: p-value *>* 0.05, ^*^ p-value ≤ 0.05, ^**^ p-value ≤ 0.01, ^***^ p-value ≤ 0.001, ^****^ p-value *<*= 0.0001). A,C: Boxplots show the median as center, the lower and upper hinges that correspond to the 25th and the 75th percentile, and whiskers that extend to the smallest and largest value no more than 1.5^*^IQR. Values that stray more than 1.5^*^IQR upwards or downwards from the whiskers are considered potential outliers and represented with dots.

**Figure S2.**
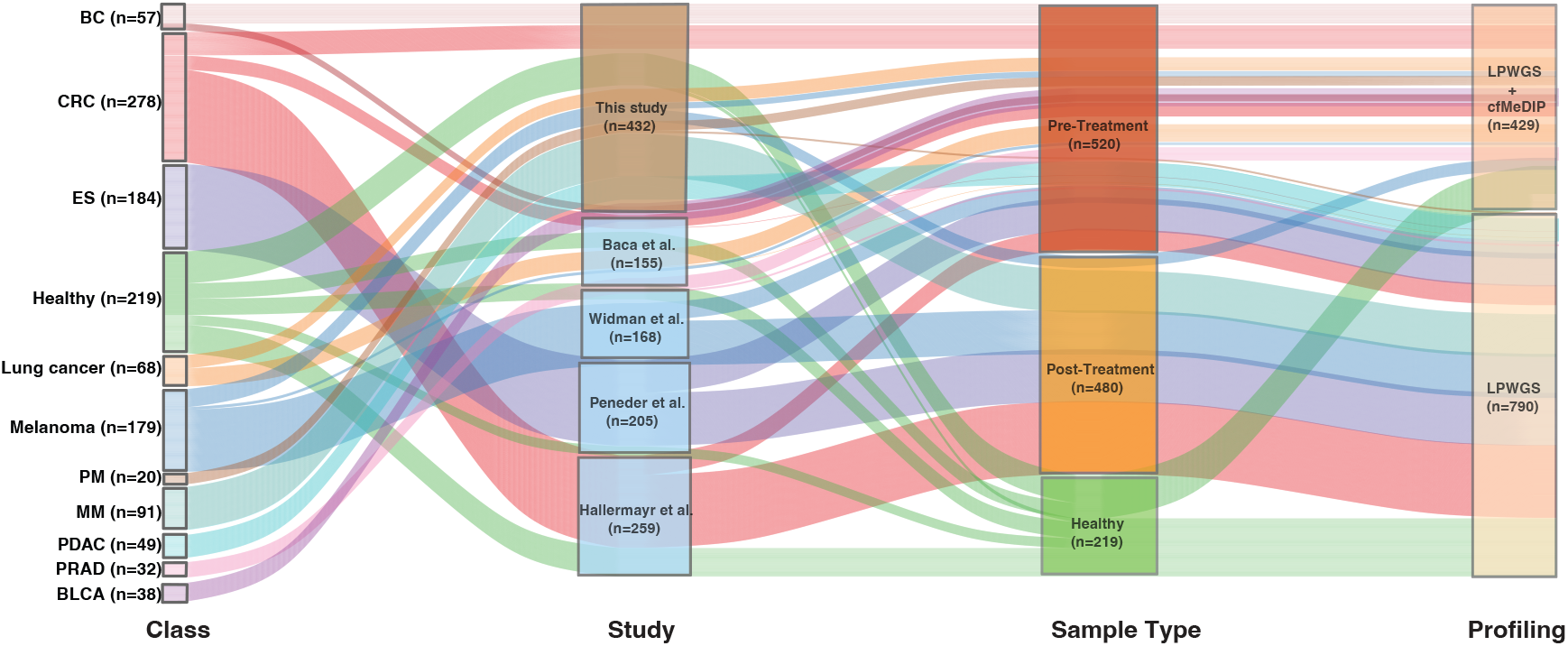
Data Cohort. Sankey diagram showing the composition of the study cohorts. Samples are stratified by class (left), study of origin (middle-left), internal data (brown), external data (light blue), sample type (middle-right), and sequencing-based approaches (right). The diagram illustrates the samples across categories, including pre-treatment, post-treatment, and control groups, as well as low-pass whole-genome sequencing (LPWGS) and LPWGS combined with cfMeDIP-seq data. Sample counts for each category are indicated in parentheses.

**Figure S3.**
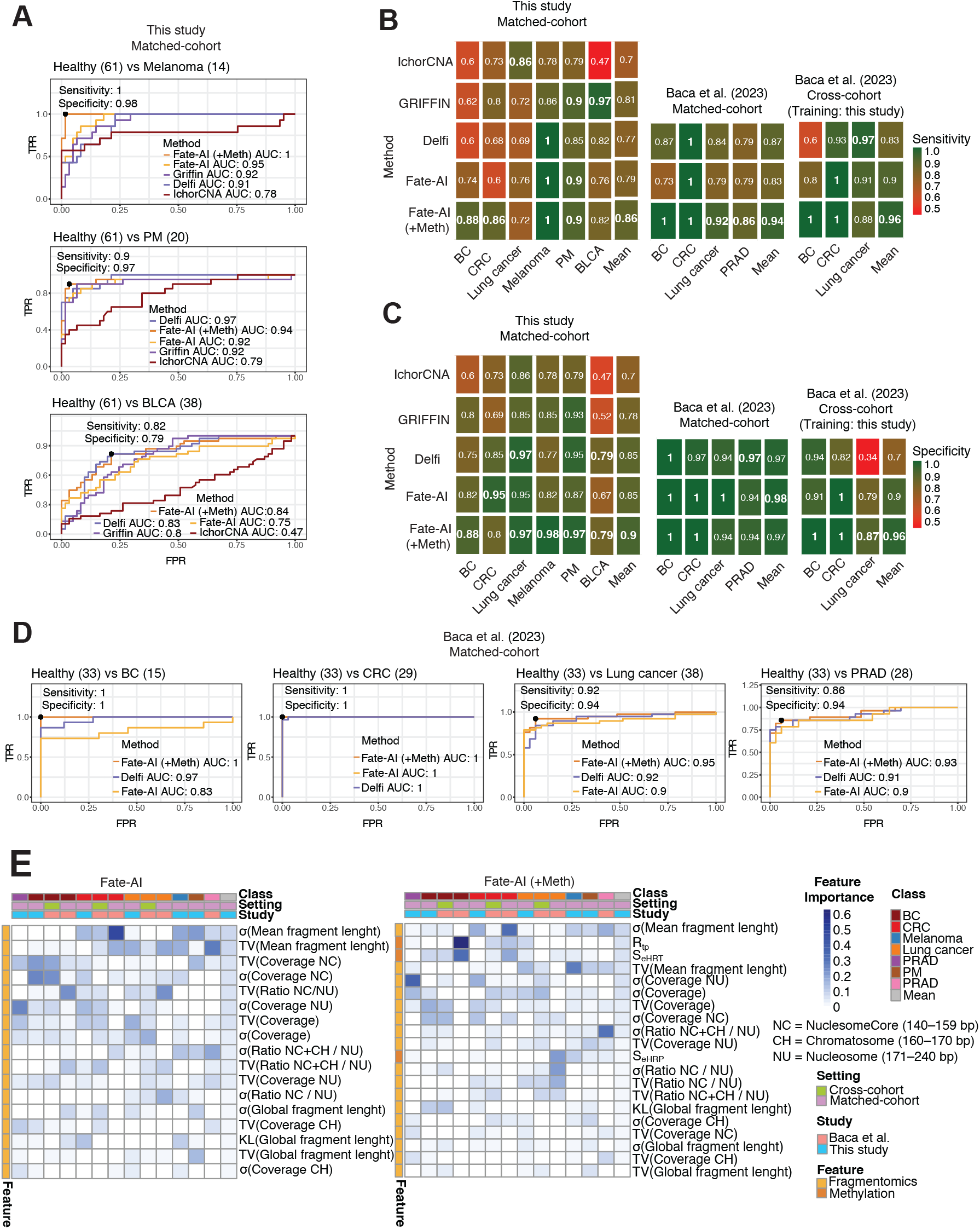
Performance evaluation of methods for cancer detection in LPWGS-cfMeDIP-seq. **A**, ROC curves illustrating classification performance in matched-cohort setting for melanoma (melanoma n=14, healthy n=61), PM (PM n=20, healthy n=61), and BLCA (BLCA n=38, healthy n=61), data from this study. **B**, Heatmap summarizing the sensitivity achieved by multiple methods in the various settings: matched-cohort on data from this study, matched-cohort on Baca et al. [25], cross-cohort on Baca et al. [25] with trained model on this study. **C**, Same as in B, for the specificity. **D**, ROC curves illustrating classification performance in matched-cohort setting for BC (BC n=15, healthy n=33), CRC (CRC n=29, healthy n=33), lung cancer (lung cancer n=38, healthy n=33), and PRAD (PRAD n=28, healthy n=33) from Baca et. al [25]. **E**, Heatmap of feature importance across different cancer types for the Fate-AI (left) and Fate-AI (+Meth) (right). Features are sorted by mean importance across studies, settings, and cancer types. Color intensity represents feature importance from low (white) to high (blue). Features are grouped into fragmentomics and methylation categories.

**Figure S4.**
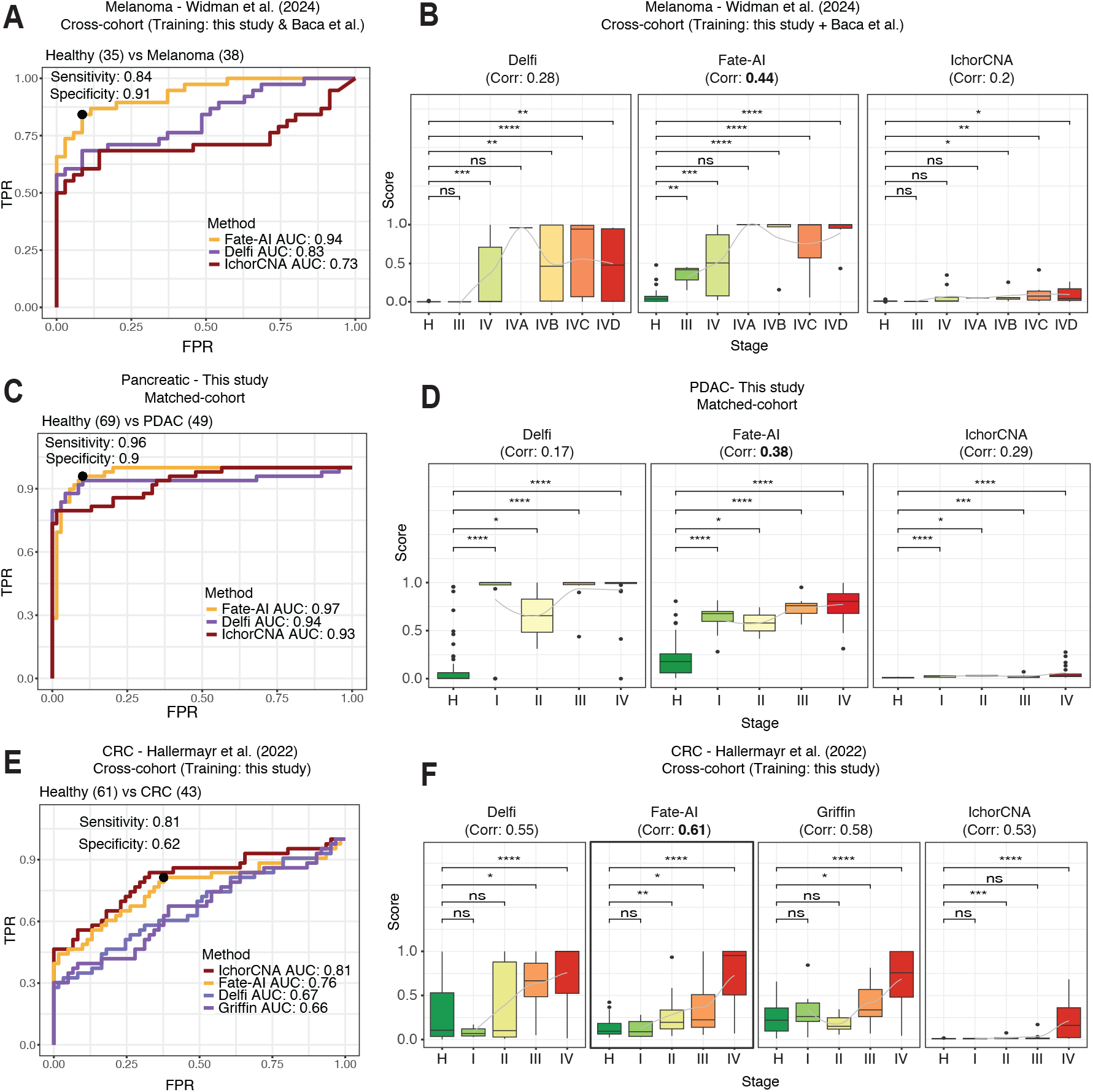
Performance evaluation of methods for cancer detection in LPWGS. **A**, ROC curves illustrating classification performance (melanoma vs healthy) when the model is trained on data from this study and Baca et. al 2023 (melanoma n=20, healthy n=94) and tested on the dataset from Widman et al. 2024 study (melanoma n=38, healthy n=35). **B**, Boxplots of predicted scores stratified by healthy (H) and cancer stage (III-IV) for melanoma (n=38). **C**, ROC curves illustrating classification performance (PDAC vs healthy) for the model trained with cross-validation with data from this study (PDAC n=49, healthy n=69). **D**, Boxplots of predicted scores stratified by healthy (H) and cancer stage (I–IV) for PDAC (n=49). **E**, ROC curves illustrating classification performance (CRC vs healthy) when the model is trained on data from this study and tested on the dataset from Hallermayr et al. study [35] (CRC n=43, healthy n=61). **F**, Boxplots of predicted scores stratified by healthy (H) and cancer stage (I–IV) for CRC (n=43). B,D,F: Boxplots represent the interquartile range (IQR), horizontal lines mark medians, and whiskers denote data points within 1.5×IQR. Significance was computed by a two−sided Wilcoxon rank-sum test (ns: p-value *>* 0.05, ^*^ p-value ≤ 0.05, ^**^ p-value ≤ 0.01, ^***^ p-value ≤ 0.001, ^****^ p-value *<*= 0.0001).

**Figure S5.**
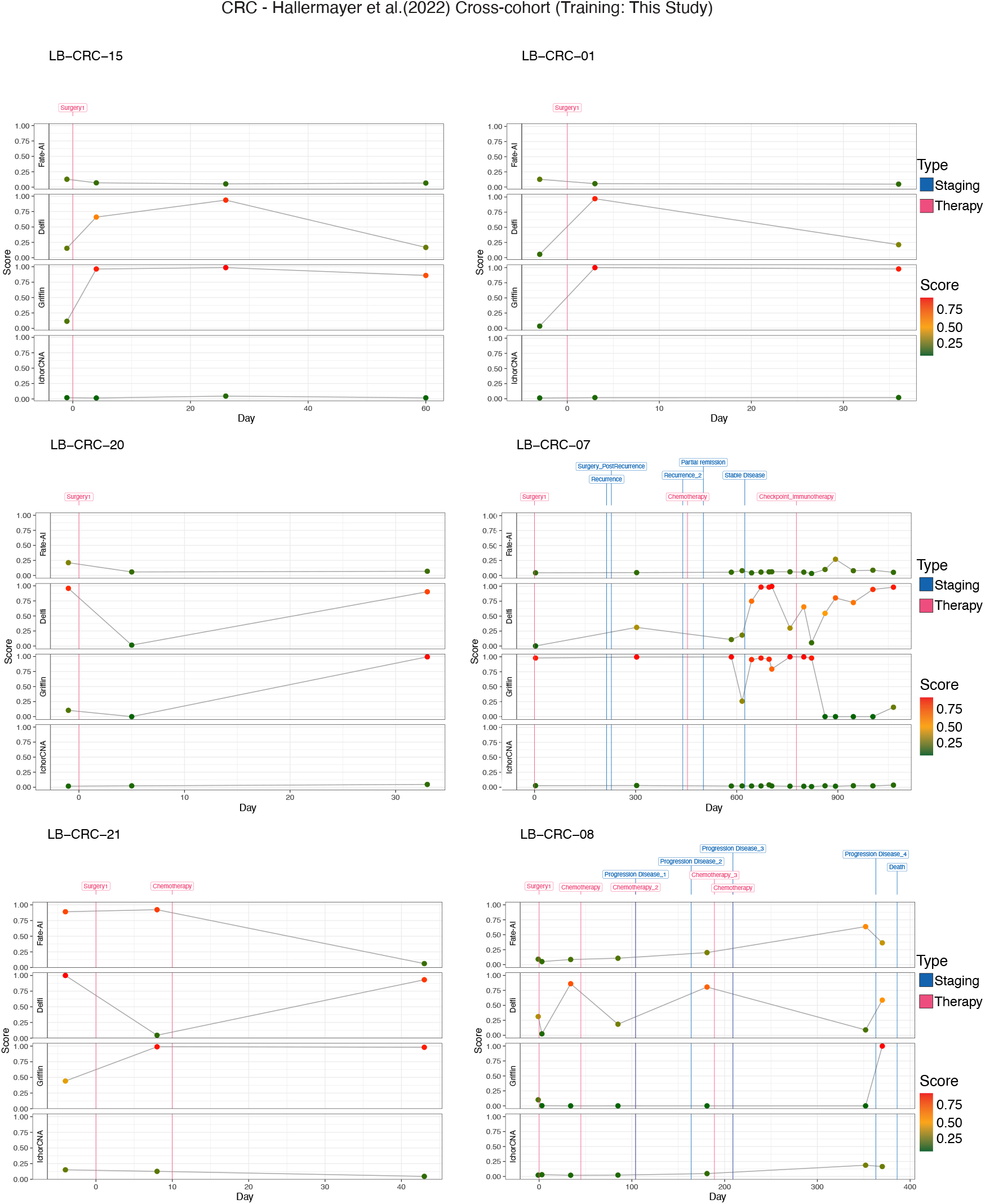
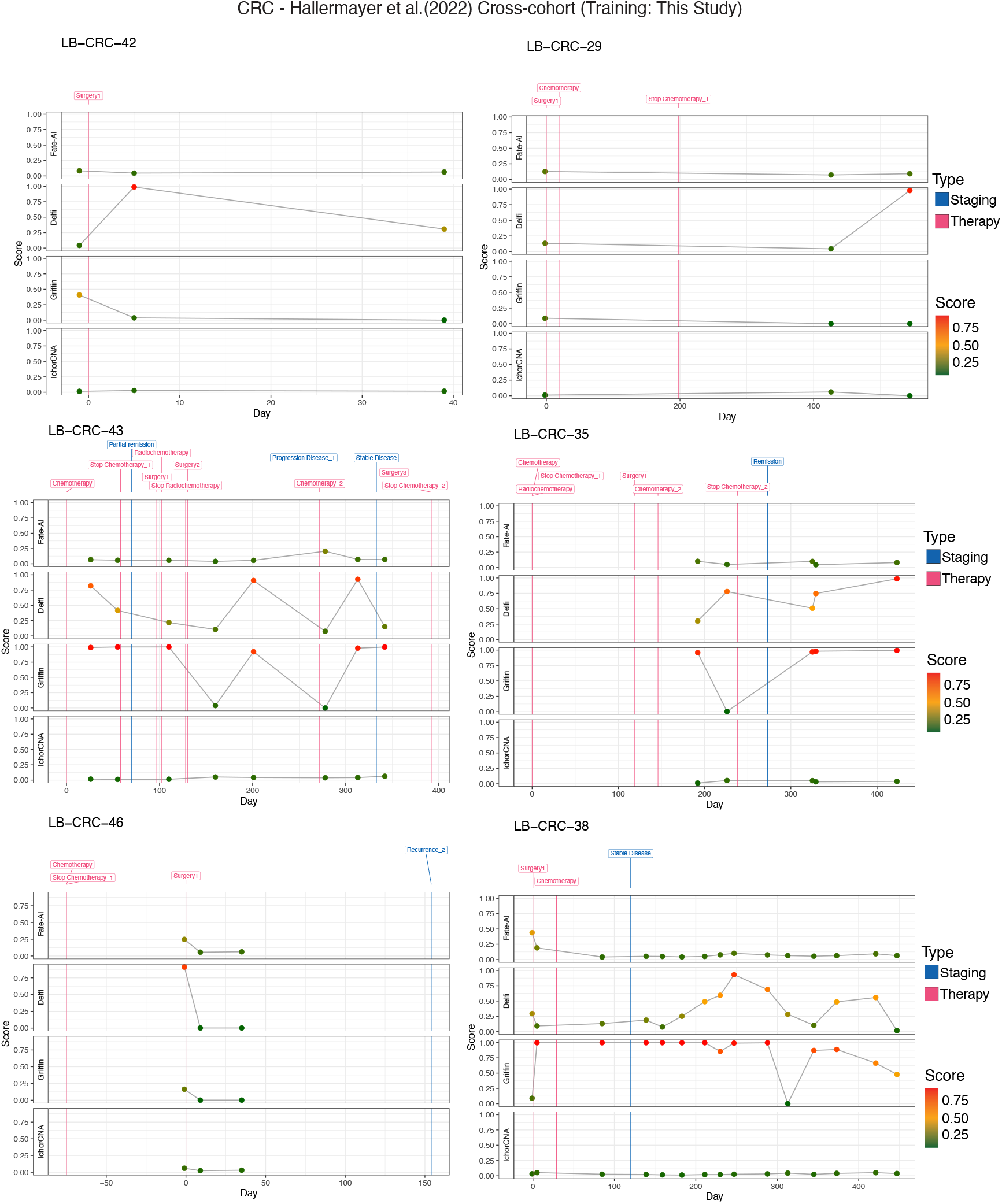
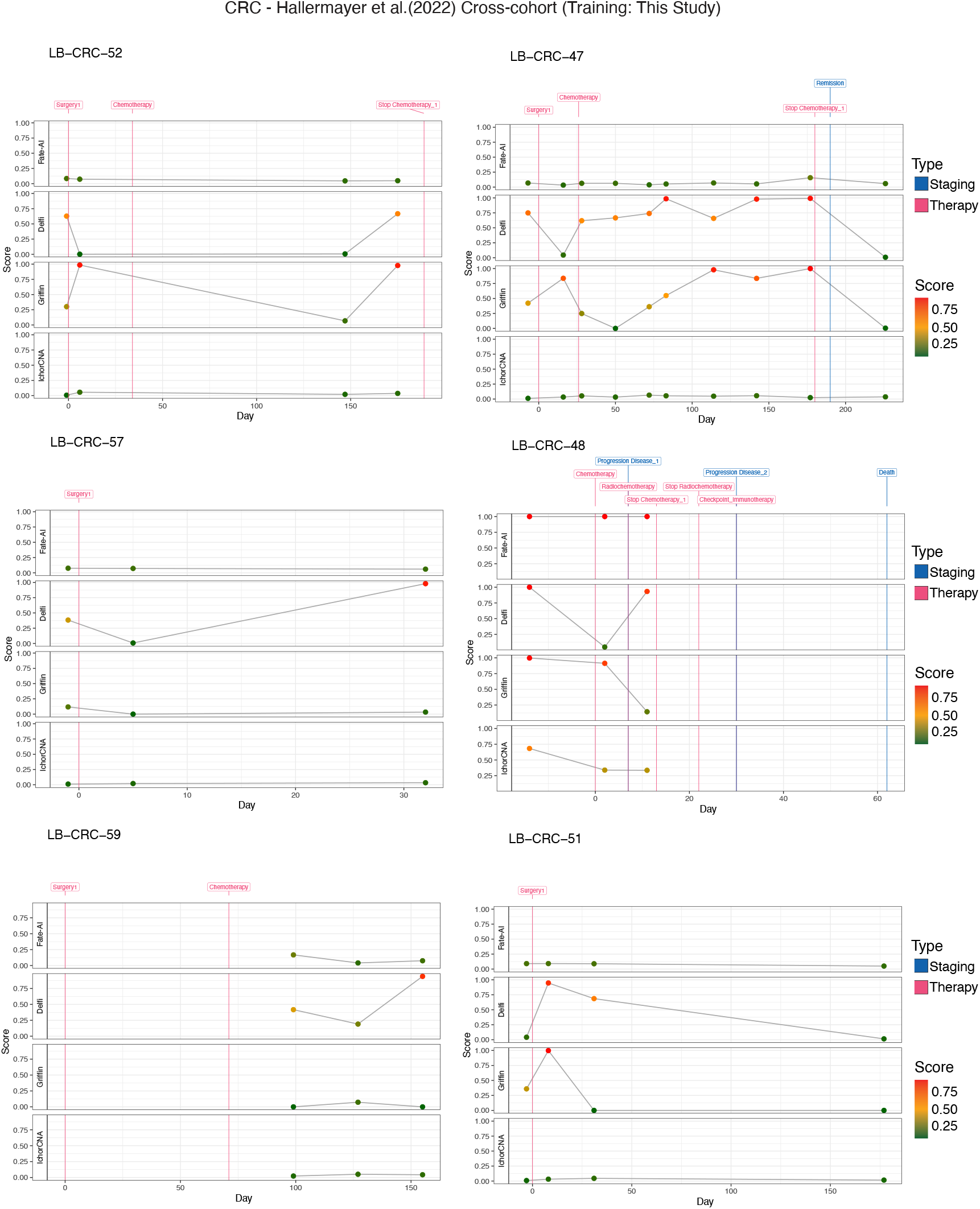
Longitudinal monitoring of colon rectal carcinoma patients. Longitudinal scores from multiple classification methods (Fate-AI, Delfi, Griffin, IchorCNA) for CRC patients (LB-CRC:1, 7, 8, 15, 20, 21, 29, 35, 38, 42, 43, 46, 51, 47, 48, 52, 57, 59) [35]. Each panel shows score dynamics over time, with therapy (red lines) and staging events (blue lines) annotated.

**Figure S6.**
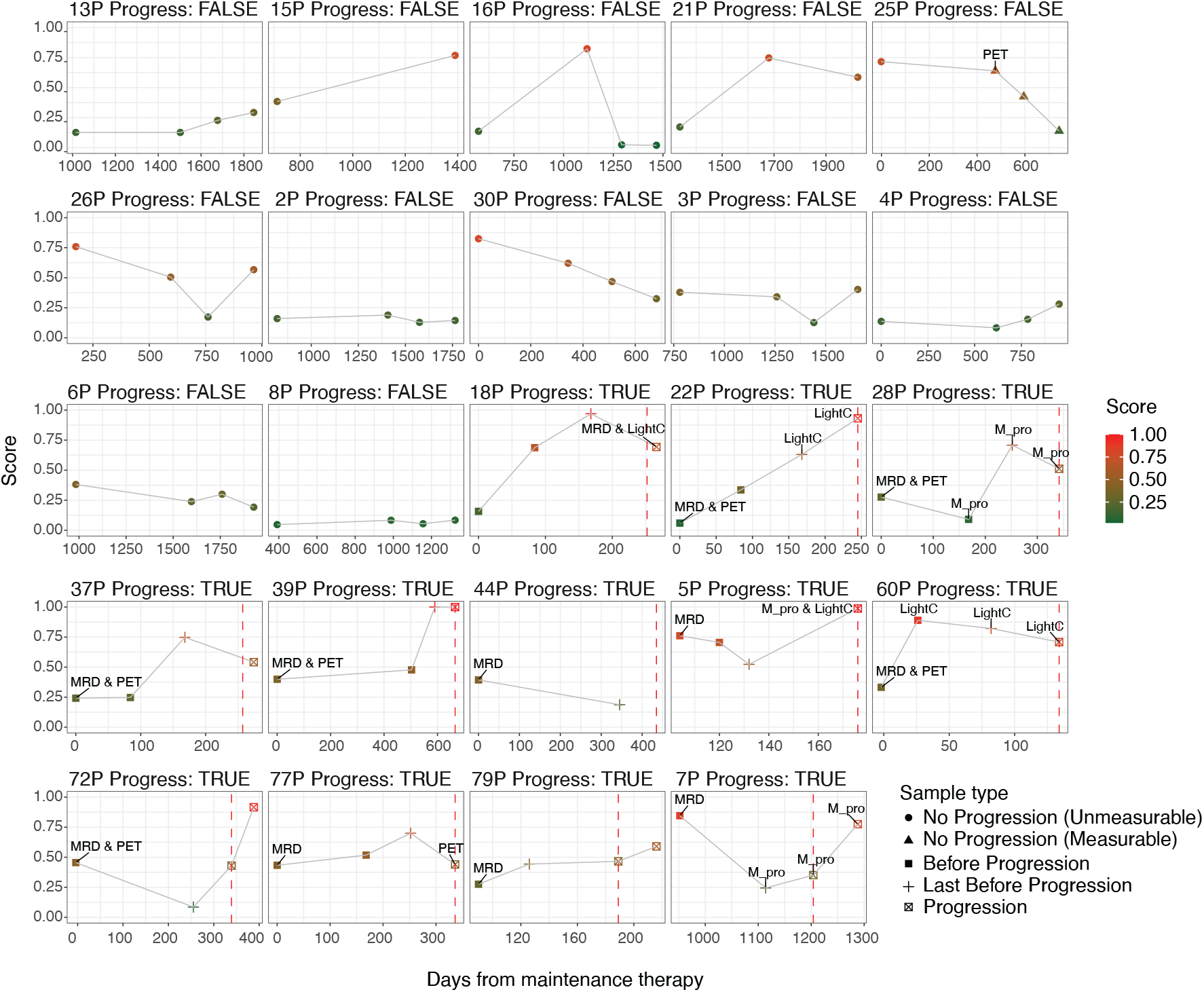
Longitudinal monitoring of multiple myeloma patients. **A**, The model was applied to 91 longitudinal samples at up to four time points (average four samples per patient, range 2-4) from 24 MM cases. Patients are stratified based on progression status: no progression (n=12) and progression (n=12). The *x* axis represents the day relative to maintenance start date and the *y* axis represents the score of Fate-AI (fragmentomic features only). Each panel shows score dynamics over time for each patient with progression event (red lines) and disease measurements that include: Monoclonal protein (M pro), Light Chain (Light C), MRD status, and Positron Emission Tomography (PET).

**Figure S7.**
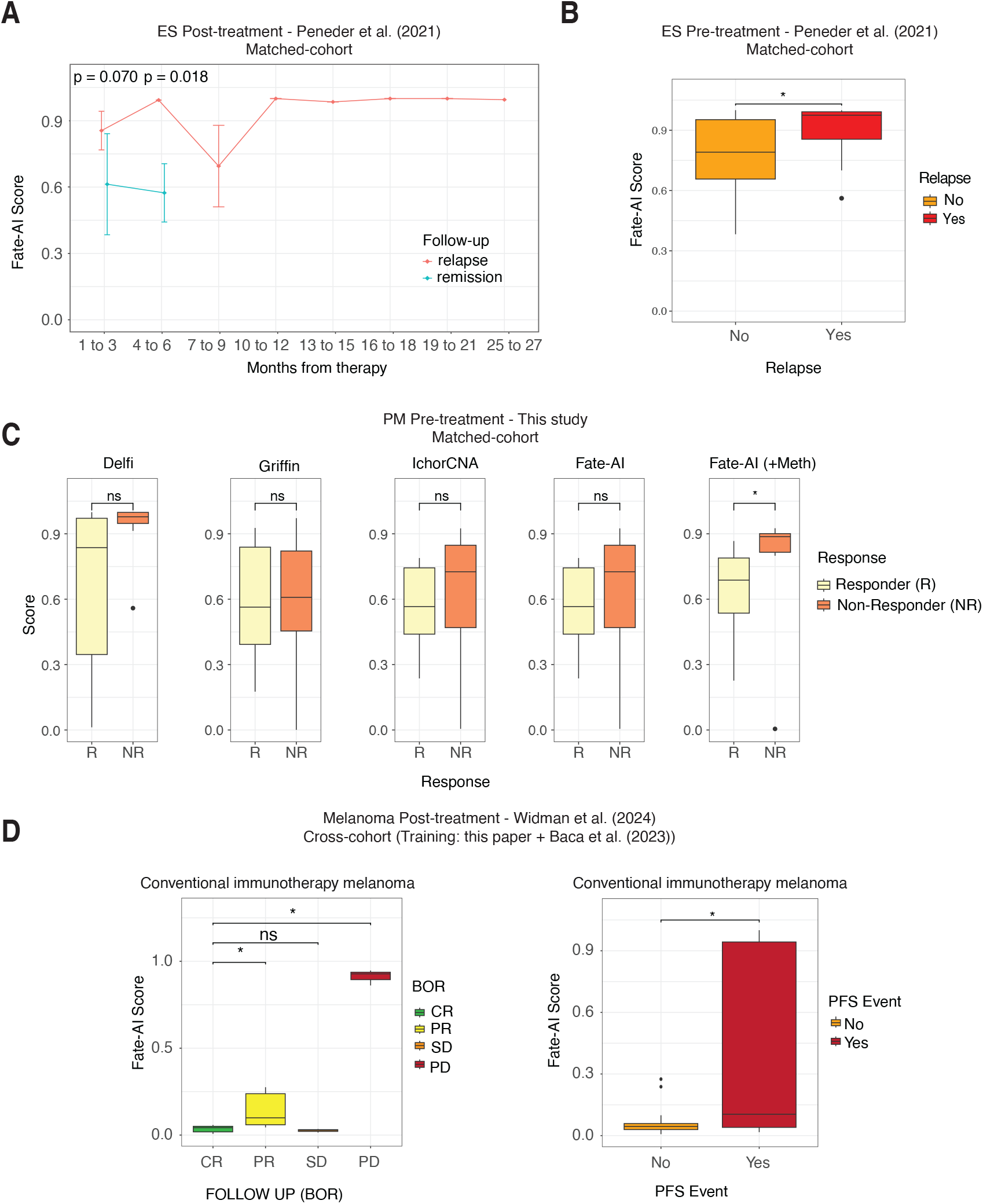
Temporal evaluation in patients before and after treatment. **A**, Post-treatment Fate-AI scores over time stratified by relapse and remission status, in the post-treatment samples of ES cohort [21]; points indicate group means and error bars represent variance. **B**, Boxplots showing Fate-AI scores (matched-cohort) between patients who experienced a relapse event and those who remained relapse-free, in the pre-treatment samples of ES cohort [21]. **C**, Scores from multiple classification methods (Fate-AI, Delfi, Griffin, IchorCNA) for pre-treatment samples of PM patients. Boxplots showing scores for each method (matched-cohort), contrasting patients with a treatment response. **D**, Boxplots showing Fate-AI scores (cross-cohort setting, training on Baca et al. [25] and this study) in post-treatment samples of the conventional immunotherapy Melanoma cohort [15]. Left panel: Fate-AI scores among BOR response categories: complete response (CR), partial response (PR), stable disease (SD), and progression disease (PD). Right panel: Fate-AI scores between patients who experienced a progression event and those who remained progression-free. B,C,D: Boxplots show the median as the center, the lower and upper hinges that correspond to the 25th and the 75th percentile, and whiskers that extend to the smallest and largest value no more than 1.5^*^IQR. Values that stray more than 1.5^*^IQR upwards or downwards from the whiskers are considered potential outliers and represented with dots. A,B,C,D: Significance was computed by a two−sided Wilcoxon rank-sum test (ns: p-value *>* 0.05, ^*^ p-value ≤ 0.05, ^**^ p-value ≤ 0.01, ^***^ p-value ≤ 0.001, ^****^ p-value *<*= 0.0001).

